# A Multi-Omics Atlas of Naturally Occurring Myelomeningocele in a Large-Animal Model Reveals Heritable Architecture

**DOI:** 10.64898/2026.05.12.724696

**Authors:** Mehmet Kizilaslan, Anas Abou Merhi, Joyce Koueik, Sydney P. Kolstad, Jamila N. Hamdan, Walid Farhat, Ligia A. Papale, Reid S. Alisch, Timothy M. George, John Wellons, Bruna Corradetti, PASTURES Consortium, Richard H. Finnell, Hasan Khatib, Bermans J. Iskandar

## Abstract

Neural tube closure relies on tightly coordinated morphogenetic programs integrating convergent extension, apical constriction of the neuroepithelium, and precise cell-cell interactions across germ layers. Disruption of these processes results in myelomeningocele, a severe complex congenital defect with lifelong multisystem consequences whose genetic and epigenetic determinants remain poorly defined. Using a sheep population with naturally occurring myelomeningocele, we quantified substantial heritability (0.42–0.68) and generated the first integrated multi-omics, multi-tissue atlas of this condition in any mammalian species. Genetic, transcriptomic, and whole-genome DNA methylation profiling across ectoderm- and mesoderm-derived tissues revealed shared and lineage-specific perturbations converging on cell-adhesion, cytoskeletal, migratory, inflammatory, and folate-responsive pathways. Chromosome 24 emerged as a multi-omics hotspot enriched for differentially expressed genes, differentially methylated regions, and candidate regulatory loci of GWAS signals, overlapping with human neurological and embryonic development trajectories. Cross-tissue network analyses highlighted coordinated disruption of neurulation-critical gene modules, establishing sheep as a robust translational model for mechanistic dissection of neural tube defect biology.

## Introduction

Vertebrate neural tube closure is a dynamic biomechanical process requiring precisely coordinated cell–cell interactions and morphogenetic movements of the neural plate synchronized with surrounding germ-layer tissues and matrix ^1,2^. Failure of neuropore fusion during neurulation leads to neural tube defects (NTDs), affecting approximately 0.3–199.4 per 10,000 births ^1,3^. Myelomeningocele (MM), or open spina bifida, arises from failed caudal neuropore closure during the third to fourth week of human gestation and leads to incomplete formation of the lower spinal cord. It is commonly accompanied by hydrocephalus and hindbrain herniation (Chiari II malformation), and associated with neurological deficits, and neurogenic bowel and bladder ^4^. Even after surgical repair, scar tissue and these comorbidities drive persistent lifelong complications that substantially compromise quality of life ^4,5^. Despite its prevalence, the underlying molecular mechanisms of MM remain poorly understood.

The current etiologic understanding of MM defines the defect as a multifactorial complex disorder of a polygenic nature, with prevalence shaped by genetic predisposition along with environmental exposures ^6–8^. Numerous epidemiological risk factors, as well as over 250 genes (MGI records) and a broad spectrum of biological pathways, have been linked to MM in mammals, resulting primarily from mouse studies and a small number of human NTD cohort studies ^6,9^. While mouse models have yielded numerous candidate NTD genes, few are over-represented in human cohorts, likely reflecting species-specific anatomical and physiological differences in neural tube development, and several genes identified in humans likewise fail to show association across mouse NTD models ^2,10–12^. These limitations highlight the need for large-animal models that better mirror human anatomy and physiology, while acknowledging that species-specific development requires dedicated MM studies for each species model.

Although MM is often cited as having a nominal heritability of 60–70%, no rigorous twin-, pedigree-, or genome-based estimate exists in any species ^6,8^. Therefore, most studies report recurrence risks or genetic association signals, lacking a robust quantitative heritability. Moreover, environmental factors, including but not limited to folate or vitamin B12 deficiency, arsenic exposure, anti-seizure medications, and maternal diabetes or obesity, also elevate MM risk ^6,8,13^. Among those, periconceptional folic acid supplementation reduces recurrence by roughly 50–70%, likely through one-carbon metabolism and *DNMT*-mediated epigenetic regulation ^6,14,15^, yet the underlying mechanisms, affected loci, and long-term consequences remain unresolved. Although widely considered safe due to its water solubility, excessive folic acid intake has been associated with adverse pregnancy outcomes, neurodevelopmental concerns, increased carcinogenic risk, and metabolic disorders, including diabetes mellitus, in both highly exposed humans and experimental mouse models ^16–18^. Understanding the epigenetic and transcriptional programs underlying MM is therefore essential for improving risk stratification, refining prevention strategies, and minimizing unintended effects of folate fortification programs.

Epigenetic mechanisms regulate gene expression beyond DNA sequence, shaping phenotypes and enabling effects that can persist across generations ^19,20^. As embryonic germ layers differentiate during neurulation, lineage-specific transcriptional programs are reinforced by environmentally responsive epigenetic modifications ^21–23^. DNA methylation marks established during this window are mitotically stable, guiding directed differentiation and propagating germ-layer-specific regulatory states into descendant tissues ^24,25^. Methylation alterations have long been implicated in NTDs, primarily through folate-mediated one-carbon metabolism but also with other environmental stimuli ^6,14^. While global methylation studies are limited, those existing studies consistently omit genetic and expression integration, lack NTD-subtype resolution, and rely heavily on single tissue mouse models with limited translational relevance and minimal validation ^26–29^. Furthermore, interactions between genetic variation and global hypomethylation secondary to folate status and pathway genes like *MTHFR* and *DNMTs*, add further complexity, reflecting the multifactorial nature of NTDs ^30,31^. The narrow scope of most previous studies limits their translational relevance, as NTDs are multi-germ layer, polygenic, and environmentally modulated traits that require integrative, genome-scale analysis.

Rapid advances in next-generation sequencing now allow simultaneous whole-genome methylation, transcriptomic, and genomic profiling from the same organisms, enabling integrative, systems biology framework analyses for complex disease risk. Although whole-genome methylation sequencing (WGMS) and RNA sequencing (RNA-Seq) have been widely applied across human and mouse disease studies, NTD research has relied almost entirely on candidate-gene methylation assays (e.g., *MTHFR, VANGL, PAX3, MGMT, GLI2*) and limited array-based methylation (∼450K CpGs), lacking any WGMS studies conducted in humans or any animal species^26,32–35^. On the other hand, genome-wide association studies (GWAS) and high-density single nucleotide polymorphism (SNP) arrays are powerful tools for mapping mammalian trait associations ^36,37^, but their application to NTDs, especially MM, remains sparse. Low incidence, inconsistent NTD phenotyping, environmental noise, and population confounders (stratification, weak linkage) have obscured the identification of strong human risk variants. Using GWAS with appropriate animal models has been deemed important for identifying genes associated with NTDs ^10^. Large animal NTDs are increasingly recognized across species, yet their incidence and heritability remain largely uncharacterized, despite developmental defects accounting for 3– 25% of neonatal and perinatal losses ^38^. Sheep offer notable translational advantages over mice, especially for naturally occurring neurological disorders, physiological alignment, and *in utero* experimental NTD repair ^39–41^. A recently identified sheep population with naturally occurring MM shows an unusually high, stable incidence (∼10% per lambing season) under controlled breeding and uniform management across decades ^42^. This population provides a rare opportunity to dissect the genetic, epigenetic, and transcriptomic architecture of MM across tissues within a systems-biology framework, while pinpointing molecular drivers that can guide prevention, management, and welfare strategies in sheep.

To date, despite the diverse genetic and environmental factors associated with MM, a holistic multi-omics analysis within a single experimental population of a large-animal model has not been reported. Our approach integrates genetic, epigenomic and transcriptomic drivers underlying disease pathology to move beyond correlative signatures and uncover candidate causal mechanisms. Considering the quantitative, polygenic and multi-layer mechanistic nature of MM, a multi-dimensional systems biology perspective is needed to unravel the causality and genetic and epigenetic architecture of this complex disorder. Thus, our investigation aimed at an integrative systems biology strategy, enabling the identification of key molecular drivers implicated in MM pathogenesis. We hypothesized that MM originates from coordinated genetic and multi-omics perturbations arising within the mesodermal and ectodermal germ layers during embryogenesis, with these early-layer disruptions subsequently propagated across the diverse tissues derived from those lineages. The objectives of our study were to (i) estimate the pedigree- and SNP-based heritability of ovine MM; (ii) examine single- and multi-omics drivers of the defect; (iii) investigate lineage-specific and cross-tissue DNA methylation and gene expression profiles associated with MM; and (iv) characterize key regulatory pathways and biologically significant hub genes.

## Results

### Strong heritable liability underlies MM risk

The study population comprised of 1,965 lambs born between 2002 and 2025, descending from 23 sires and 357 dams; 54 ewes and 4 rams formed the pedigree founder base with no recorded parents. Of these lambs, 973 were females and 987 males, with sex unrecorded for five. Birth types included 427 singletons, 1,220 twins, and 267 multiples. A total of 138 lambs (55 females, 83 males) were affected with MM, while 1,824 were unaffected and three lacked phenotype records. A generalized linear model indicated moderate evidence of a sex effect, with males showing significantly higher odds of MM (p < 0.05), corresponding to a 2.8-percentage-point increase in predicted risk (95% CI: 0.5–5.1%). MM-affected lambs also had significantly lower birth weight (–0.66 ± 0.21 lb; p < 0.05). The population exhibited high relatedness, with a mean inbreeding of 0.15 ± 0.002 and an effective population size of ∼76.

To estimate liability-scale heritability (*h*^*2*^) for MM, we used a Bayesian approach based on pedigree data from 2002 to 2025. We identified a moderate-to-high *h*^2^ for MM, with a posterior mean of 0.68 (95% CI: 0.40-0.90). On the other hand, the Bayesian estimation of SNP-based genomic *h*^*2*^ for MM was 0.42 (95% CI: 0.14-0.85), with a larger yet corresponding confidence interval than the pedigree-based heritability. Both estimates suggest a moderate to strong liability to MM due to genetic factors.

### GWAS reveals a 5-Mb chromosome 24 interval with multiple potential QTL

As part of a more targeted effort to define the genetic risk factors contributing to MM, we performed a Principal Component Analysis (PCA)-adjusted GWAS including 88 cases and 100 controls. This analysis identified a genomic interval on ovine chromosome 24 (Chr24) significantly associated with MM (**Fig. 1A; table S1**). In total, 13 genome-wide significant SNPs mapped to Chr24, spanning the 25-30Mb region, led by an intronic SNP in Integrin Subunit Alpha D (*ITGAD*) with an odds ratio of 1.04 (4.06% increased risk per allele). Another genome-wide significant SNP was found within the 5^th^ exon of TLC domain containing 3A (*TLCD3A*) gene with a 4.80% change in odds, and four SNPs on chromosomes 16, 22, 2 and 3 showed suggestive effects (2.41–11.38%; **table S1**).

**Figure 1.**
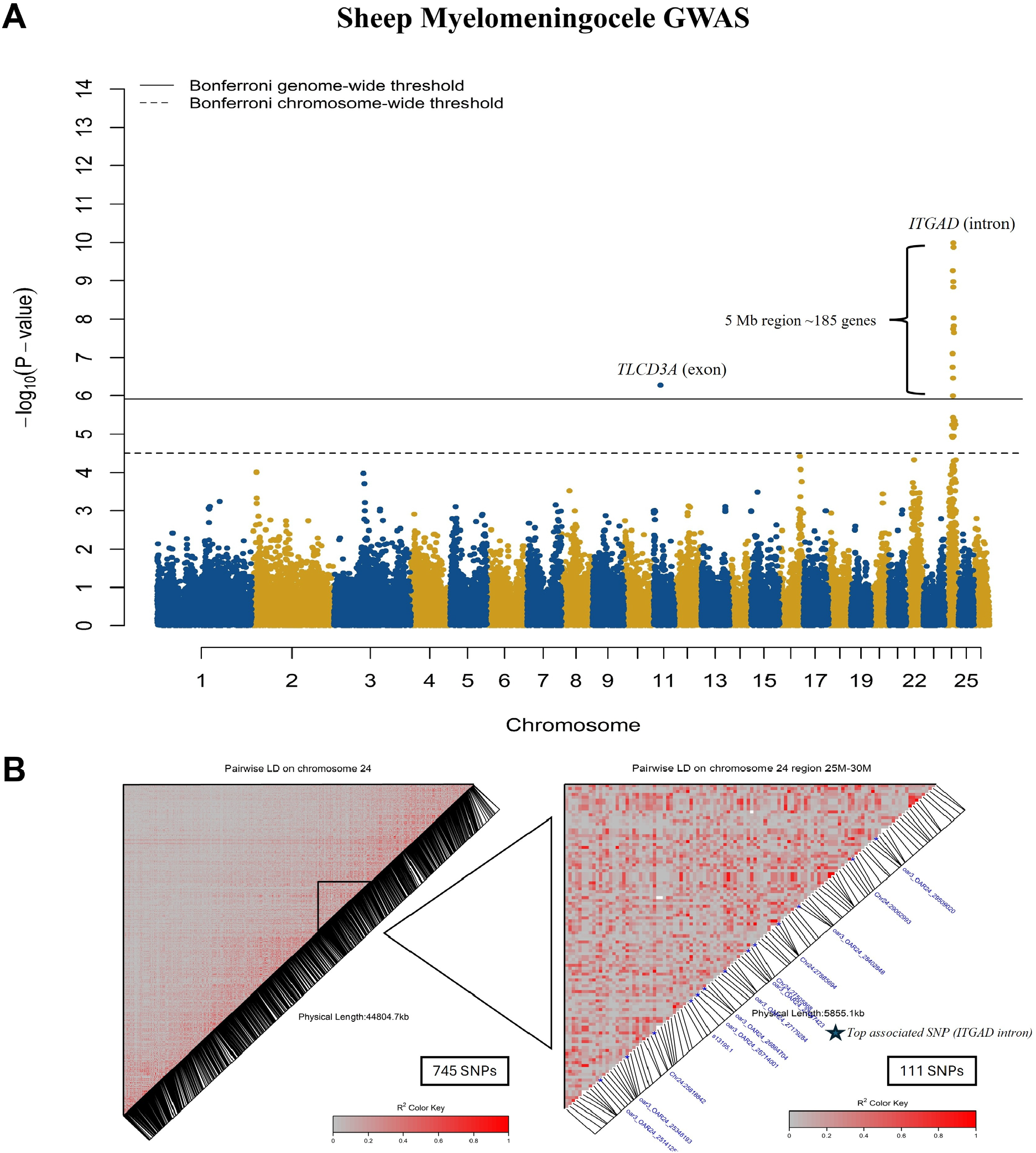
Manhattan plot of GWAS for MM and LD heatmap on Chr24. **(A)** Manhattan plot of GWA analysis is presented with SNPs distributed across 26 sheep autosomes on the x-axis and their associated p-values given in - log10 scale on the y-axis. Solid black line represents a Bonferroni-corrected genome-wide significance threshold and dashed chromosome-wide significance threshold. Points are SNPs scattered across chromosomes (x-axis) and association signal (y-axis) with MM. **(B)** Pairwise LD heatmap of the Chr24 (left) and associated genomic interval (right) with MM. Based on the scale presented in the legend, gray represents no LD, red represents complete LD, and the area between gray and red shows pairwise LD (*r*^*2*^) distributed between 0 and 1. The star on the heatmap (right) marks the location of top associated SNP in *ITGAD*.

Linkage disequilibrium (LD) analysis on Chr24 (**Fig. 1B**) showed pairwise squared correlation coefficients (*r*^*2*^) among the 13 significant SNPs ranging from 0.18 to 0.95, indicating moderate to high correlation. Extensive long-range LD between SNPs, consistent with small effective population size and high relatedness, indicates that association signals span broad genomic intervals rather than resolving to a single causal location. We therefore identified multiple candidate genes per genome-wide significant locus. A conditional GWAS, incorporating the lead SNP as a covariate, abolished the association signal for the remaining 12 Chr24 SNPs. However, given the limited sample size and the extended LD structure, these results do not provide conclusive evidence against multiple QTL in the region.

Functional interrogation of the associated 5 Mb interval identified 185 genes within this relatively compact genomic region, including 128 annotated genes and 57 predicted loci (**table S2**). Notably, this gene-dense region contains several clustered gene families, many of which harbor genome-wide associated SNPs. Collectively, the highly inbred background, modest sample size, extensive long-range LD, the large 5 Mb span of the interval, and its gene-rich functionally clustered architecture indicate that this interval is biologically complex and likely harbors multiple candidate genes contributing to the observed association.

Our data further indicates that the association signal in this 5 Mb region is largely captured by the lead SNP in *ITGAD*, making it the most compelling candidate gene at this location. Nonetheless, the presence of multiple functionally related genes (including *ITGAD, ITGAM, ITGAX, TYW1, PRSS53, PRSS8, PRSS36*, etc.) and regulatory elements (e.g., *ZNF646, ZNF668, ZNF629*) with the LD structure suggests that the underlying architecture may involve more than one functional variant. Accordingly, while we propose *ITGAD* as the primary positional candidate, the broader interval may still harbor additional variants and genes contributing to MM occurrence in this sheep population. Although it has a strong association with MM, the SNP located in the exon of *TLCD3A* (**table S1**) was excluded from further analysis due to positional discrepancies between the two sheep reference genome assemblies. This variant will be reconsidered once its genomic position is resolved in future assembly updates.

### Whole-genome DNA methylation profiling reveals tissue- and germ layer–specific epigenetic dysregulation in MM

Following the identification of the associated genomic region on Chr24, we sought to characterize epigenetic signatures of MM using WGMS methylation profiling. We conducted tissue-specific single-base resolution differential methylation analysis for muscle (mesoderm), skin (mesoderm + non-neural ectoderm), and spinal cord (neuro-ectoderm) sampled adjacent to the site of the lesion in MM lambs vs. half-sib controls (CT). Of the 23,071,167 CpG cytosines tested in muscle, 4,179 hyper- and 3,911 hypo-differentially methylated cytosines (DMCs) were identified in MM lambs (**Fig. 2A, table S3**). In skin, 3,666 hyper- and 3,538 hypomethylated DMCs were detected among 18,612,656 CpG sites (**Fig. 2B, table S3**). In spinal cord, 2,590 hyper- and 1,442 hypomethylated DMCs were identified from 21,905,030 CpG cytosines (**Fig. 2C, table S3**). Converging with the GWAS results, Chr24 exhibited the highest density of DMCs in both muscle and spinal cord and remained among the most enriched chromosomes in skin (**Fig. 2D, E, F)**.

**Figure 2.**
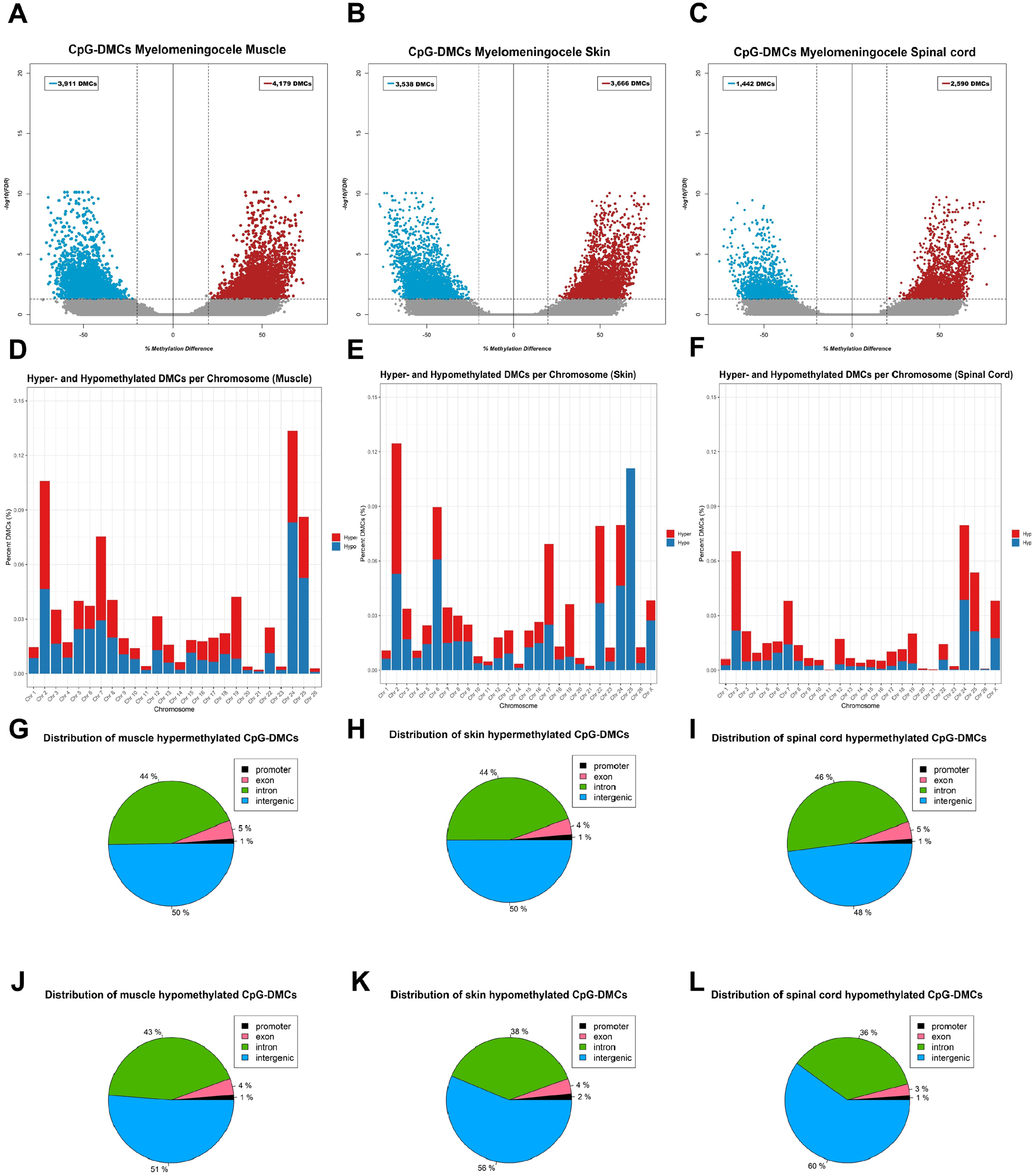
Distributions of CpG-DMCs identified in muscle, skin, and spinal cord tissues of MM vs. CT lambs. **(A)** Volcano plot of CpG-DMCs observed for muscle; **(B)** Volcano plot of CpG-DMCs observed for skin; **(C)** Volcano plot of CpG-DMCs observed for spinal cord; **(D)** Percentage of hyper- and hypomethylated DMCs in muscle per each chromosome; **(E)** Percentage of hyper- and hypomethylated DMCs in skin per each chromosome **(F)** Percentage of hyper- and hypomethylated DMCs in spinal cord per each chromosome; **(G)** Genomic context annotation of hypermethylated muscle DMCs; **(H)** Genomic context annotation of hypermethylated skin DMCs; **(I)** Genomic context annotation of hypermethylated spinal cord DMCs; **(J)** Genomic context annotation of hypomethylated muscle DMCs; **(K)** Genomic context annotation of hypomethylated skin DMCs; **(L)** Genomic context annotation of hypomethylated muscle DMCs.

Across all three tissues, most CpG-DMCs mapped to intronic or intergenic regions. In muscle, hypermethylated DMCs were distributed as 44% intronic, 5% exonic, 1% promoter, and 50% intergenic, while hypomethylated sites were 43%, 4%, 1%, and 52%, respectively (**Fig. 2G, J**). Skin showed a similar pattern, with hypermethylated DMCs at 44% intronic, 4% exonic, 1% promoter, and 51% intergenic, and hypomethylated DMCs at 38%, 4%, 2%, and 56% (**Fig. 2H, K**). In spinal cord, hypermethylated DMCs were 46% intronic, 5% exonic, 1% promoter, and 48% intergenic, whereas hypomethylated DMCs were 36%, 3%, 1%, and 60%, respectively (**Fig. 2I, L**).

CpG-DMC annotation identified 1,670 differentially methylated genes (DMGs) in muscle, comprising 613 exclusively hypermethylated, 739 exclusively hypomethylated, and 318 that contained both hyper- and hypomethylated sites. In skin, 1,448 DMGs were identified, including 593 exclusively hypermethylated, 575 exclusively hypomethylated, and 280 that contained both types of DMCs. In spinal cord, 840 DMGs were detected, comprising 491 exclusively hypermethylated, 255 exclusively hypomethylated, and 94 with both hyper- and hypomethylated DMCs. All DMGs for each tissue are listed in **table S4**.

Intersection of DMGs with concordant hyper- or hypomethylation across mesodermal and ectodermal tissues revealed that most DMGs were tissue-specific, although a substantial subset was shared among tissues (**Fig. 3**). Muscle and skin shared 802 MM-associated DMGs (402 hypermethylated and 400 hypomethylated), while 868 genes were unique to muscle and 646 to skin. In the muscle-spinal cord comparison, 566 DMGs were shared (378 hyper- and 188 hypomethylated), with 1,114 muscle-specific and 274 spinal cord-specific genes. Similarly, skin and spinal cord shared 462 DMGs (312 hyper- and 150 hypomethylated), with 986 and 378 tissue-specific genes, respectively. Across all tissues, 395 DMGs (263 hyper- and 132 hypomethylated) DMGs were consistently associated with MM (**Fig. 3**).

**Figure 3.**
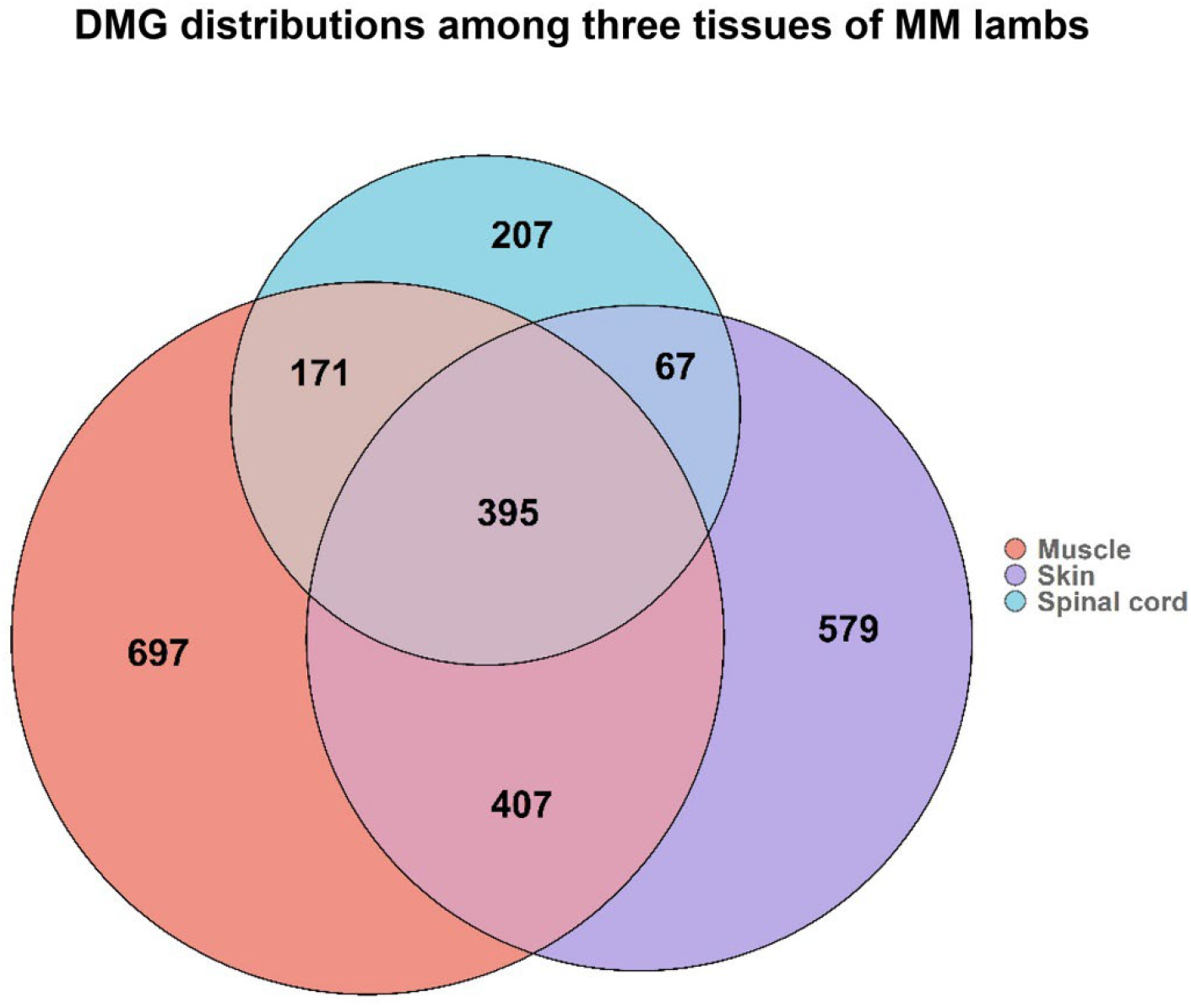
Distribution of DMGs found by differential methylation analysis of each tissue -muscle, skin, and spinal cord.

### RNA-seq profiling identifies lineage-dependent gene expression alterations in MM

To identify gene expression alterations associated with MM, we performed differential expression analysis with RNA-Seq profiles of muscle, skin, and spinal cord of MM and CT lambs. As shown in **Fig. 4A**, PCA separated the three tissue transcriptomes primarily by tissue type rather than disease state. Skin samples clustered between muscle and spinal cord, consistent with their mixed developmental origin. The first two principal components accounted for 72.5% of the total variance in gene expression. Differential expression analysis identified 1,252 differentially expressed genes (DEGs) in muscle, including 708 upregulated and 544 downregulated in MM lambs (**Fig. 4B**). In skin, 315 DEGs were detected (58 upregulated, 257 downregulated) (**Fig. 4C**), whereas spinal cord exhibited 94 DEGs (13 upregulated, 81 downregulated) (**Fig. 4D**). Complete DEG lists for each tissue are provided in **table S5**.

**Figure 4.**
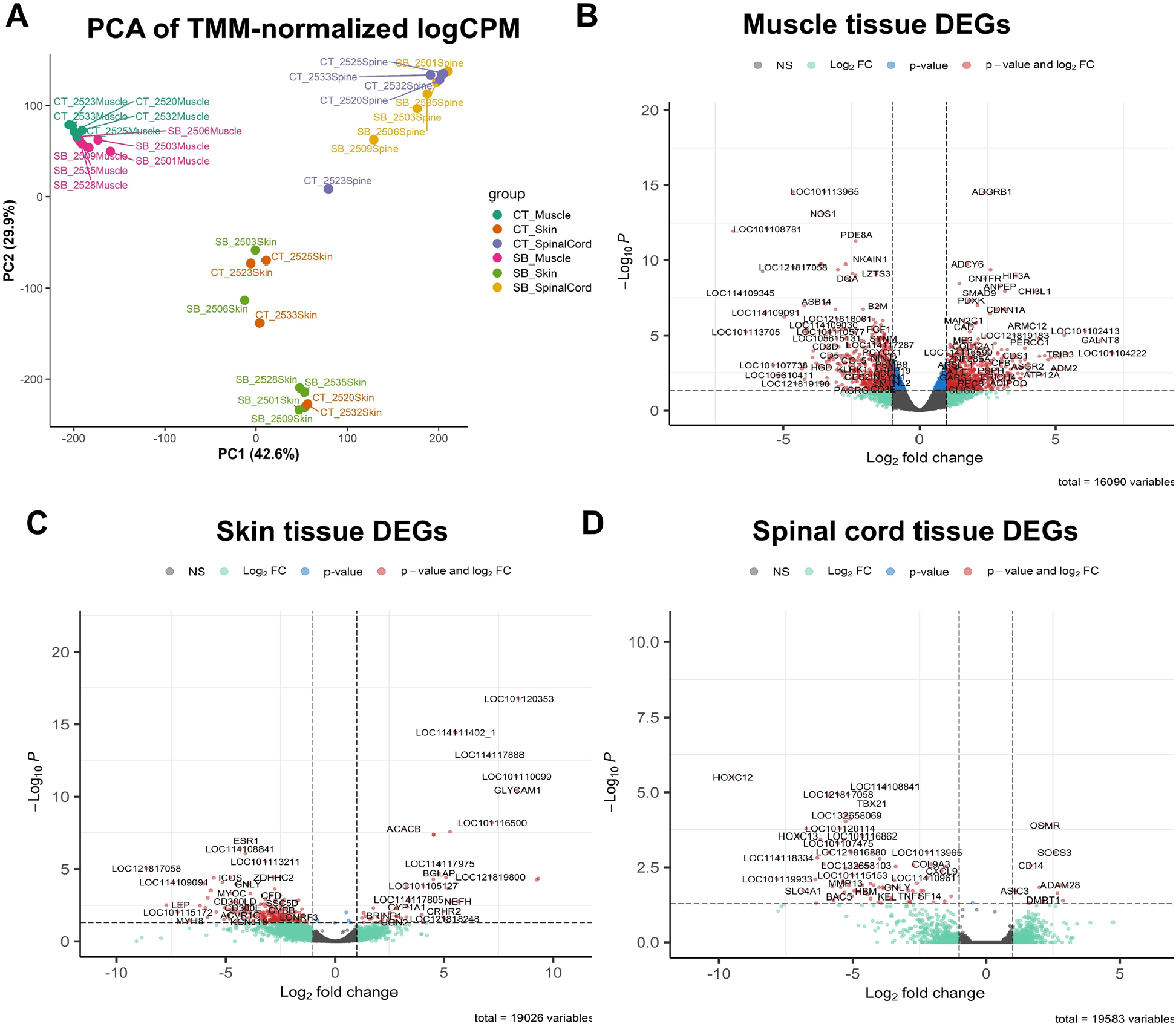
PCA and volcano plots of muscle, skin, and spinal cord tissues showing altered expression in MM vs. CT lambs. **(A)** PCA of the TMM-normalized RNA-Seq read counts on log (Counts Per Million) scale, showing clear groups of tissues with muscle having separation between MM vs. CT groups while the other two tissues are widely spread in terms of expression levels suggesting wide dispersion within MM and CT groups; **(B)** Volcano plot for muscle DEGs; **(C)** Volcano plot for skin DEGs; **(D)** Volcano plot for spinal cord DEGs. Vertical dashed lines are fold change (FC) > 2 thresholds on log2 scale, horizontal line is q < 0.05 threshold on -log10 scale, while each dot with colors represents expression level differences of genes between MM vs. CT lambs, according to their log2 fold-change (x-axis) and FDR corrected p-values (y-axis).

Pairwise comparisons showed that muscle and skin shared the highest number of DEGs in MM lambs (58), followed by muscle-spinal cord (23) and skin-spinal cord (18), with 11 DEGs common to all three tissues (**Table 1**).

**Table 1.** Pairwise DEG overlaps between tissues of different embryonic lineages.

| Tissues | Number of Shared DEGs | Gene Official Symbols |
| --- | --- | --- |
| Muscle-Skin | 58 | *ACACB, CRHR2, CCER2, CD74, OVAR-DRB1, DQA, GNLY, RGS5, CD3D, IKZF3, ESR1, TMOD2, CD2, CXCR6, GIMAP1, KLRK1, TLR7, NTN4, GIMAP6, PTPRC, CD96, LYZ, CD3E, CD300E, CD27, TNFSF8 + 32 LOC genes* |
| Muscle-Spinal cord | 23 | *FKBP5, OSMR, DMBT1, GNLY, CXCR6, RERGL, CXCL9, CD96 + 15 LOC genes* |
| Skin-Spinal cord | 18 | *GNLY, TNFSF14, TBX21, C1QTNF3, SH2D1A, CXCR6, CD96, + 11 LOC genes* |
| Muscle-Skin-Spinal cord | 11 | *GNLY, CXCR6, CD96 + 8 LOC genes* |

### Integrating GWAS with DNA methylation alterations implicates potential genome-methylome interactions underlying MM

We intersected the genome-wide significant interval on ovine Chr24 with DMGs in each tissue of MM lambs and found a striking overlap between genes implicated by genomic variation and those exhibiting epigenomic alterations, suggesting a potential interplay underlying disease etiology. Of the 185 characterized and uncharacterized genes in this region, 44 genes were shared between the GWAS and the methylation analyses in the MM muscle. Comparable overlaps were observed in skin (32 genes) and spinal cord (16 genes) (**table S6**). Of the 44 muscle DMGs, 16 (*ITGAD, ZG16, PRSS53, PRSS36, C24H16orf54, TMEM265, VKORC1L1, GUSB, ASL, CRCP, KATNIP, GSG1L,XPO6, CALN1, GALNT17*, and *AUTS2*) directly overlapped with positional candidate genes for the genome-wide significant (**table S1)**, whereas the remaining 28 DMGs were located within the same 5 Mb region on Chr24. In skin, 12 genes (*ZNF629, PRSS53, KATNIP, GSG1L, XPO6, ZG16, STX4, VKORC1L1, ASL, CALN1*, and *GALNT17*) overlapped between GWAS and methylation analyses, while 7 genes (*KATNIP, GSG1L, SBK1, PRSS36, ITGAD, CALN1*, and *GALNT17*) were shared in spinal cord. Furthermore, among suggestive SNPs identified on chromosomes 2, 3, 16 and 22 (**table S1**), *XDH* was hypermethylated in muscle and *SORCS3* was hypomethylated in spinal cord.

### GWAS-DEG integrations imply a multi-tissue link between genetic association and altered expression underlying MM etiology

To investigate potential interactions between genetic loci associated with MM and transcriptional changes, we intersected genes within the 5 Mb GWAS-significant interval on Chr24 with DEGs identified across tissues of MM lambs. This analysis yielded eight muscle DEGs (*CORO1A, LOC105604728, SPN, HSD3B7, ZNF48, DCTPP1, PRSS53*, and *PSPH*) and two skin DEGs (*ITGAL* and *DOC2A*) that overlapped with genes in the 5 Mb GWAS interval.

Motivated by the overlap of 10 genes between the two distinct analyses and considering the conservative GWAS threshold, small sample size, and strong overdispersion correction in the skin and spinal cord DEG datasets, we expanded the evaluation to include genes showing at least a 2-fold expression change without enforcing the statistical significance cutoff (q < 0.05). Under this relaxed criterion, only one additional gene in muscle (*GDPD3*) overlapped with the genomic interval on Chr24. In contrast, the number of genes with altered expression within the region increased substantially in skin, rising from two to 16 (*ITGAL, DOC2A, ITGAM, CORO1A, CD19, SPN, LAT, ATP2A1, CALN1, GUSB, APOBR, HSD3B7, PHKG1, CTF1, TPST1* and *PRSS53*), and to seven in spinal cord (*ITGAD, ITGAM, LAT, SPN, PRRT2, STX1B*, and *AHSP*). Among these genes, *SPN* overlapped across all three tissues, while *PRSS53, HSD3B7* and *CORO1A* were shared between muscle and skin, and *LAT* and *ITGAM* overlapped between skin and spinal cord. Notably, several of these genes (*ITGAD, ITGAM, PRSS53, LAT* and *SPN)* are also positional candidate genes reported in the GWAS analysis (**table S1, S5)**.

### Integration of DMGs and DEGs highlights altered methylation as a potential driver of dysregulated gene expression in MM

To identify genes whose expression may be regulated by DNA methylation changes, we intersected DMGs with DEGs identified in each tissue of the MM lambs. This analysis revealed multiple genes showing concurrent alterations in both methylation and expression across tissues. Specifically, 89 genes in muscle exhibited both differential methylation and expression in MM lambs compared with 14 in skin and 6 in spinal cord (**Table 2, table S7**).

**Table 2.**
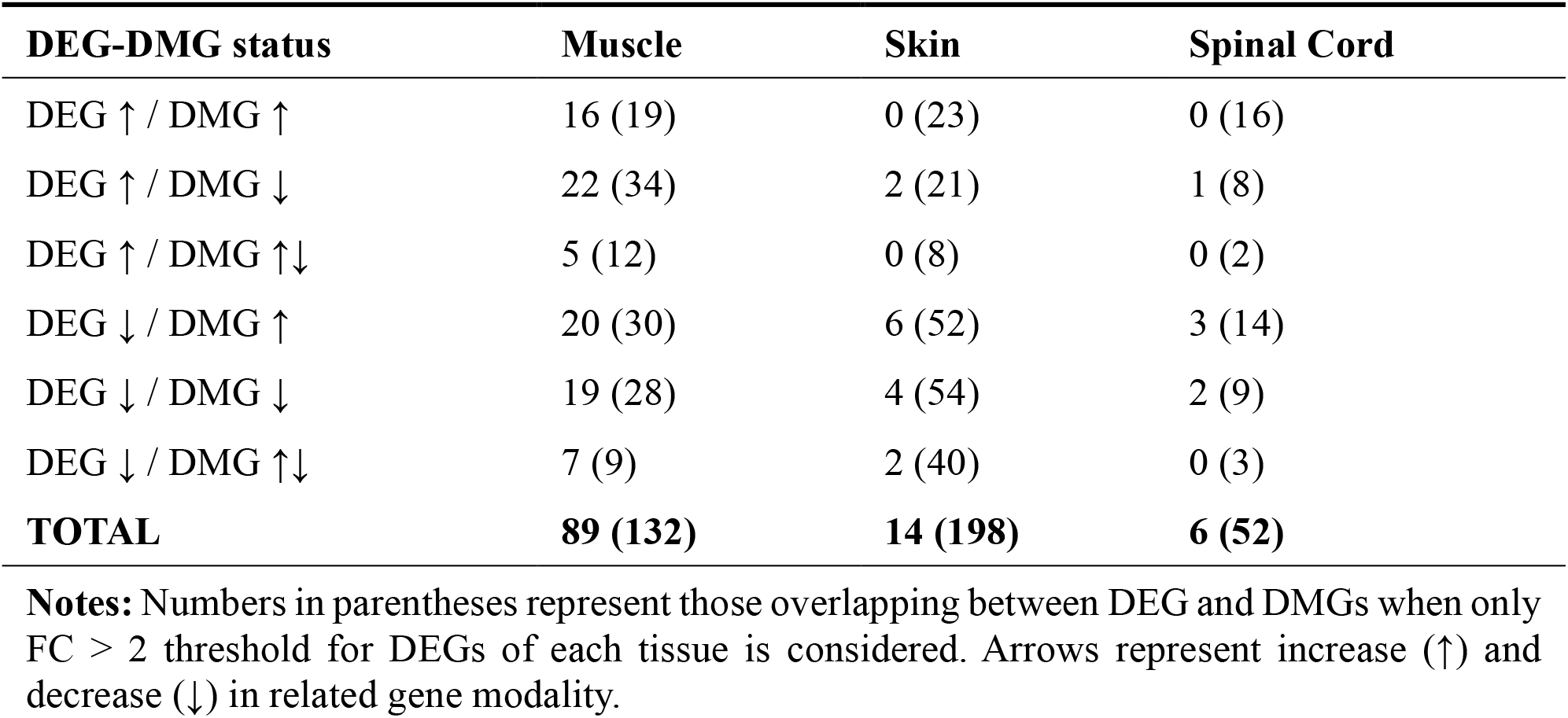
Intersection of DMGs and DEGs of muscle, skin, and spinal cord in MM lambs.

| DEG-DMG status | Muscle | Skin | Spinal Cord |
| --- | --- | --- | --- |
| DEG ↑ / DMG ↑ | 16 (19) | 0 (23) | 0 (16) |
| DEG ↑ / DMG ↓ | 22 (34) | 2 (21) | 1 (8) |
| DEG ↑ / DMG ↑↓ | 5 (12) | 0 (8) | 0 (2) |
| DEG ↓ / DMG ↑ | 20 (30) | 6 (52) | 3 (14) |
| DEG ↓ / DMG ↓ | 19 (28) | 4 (54) | 2 (9) |
| DEG ↓ / DMG ↑↓ | 7 (9) | 2 (40) | 0 (3) |
| **TOTAL** | **89 (132)** | **14 (198)** | **6 (52)** |

**Notes:** Numbers in parentheses represent those overlapping between DEG and DMGs when only FC > 2 threshold for DEGs of each tissue is considered. Arrows represent increase (↑) and decrease (↓) in related gene modality.

To further assess the relationship between methylation and transcriptional changes, we expanded the analysis to include genes exhibiting at least 2-fold expression difference without applying the statistical significance threshold (q < 0.05). Under this criterion, the number of genes showing concurrent expression and methylation changes increased modestly in muscle to 132 genes. In contrast, substantially larger increases were observed in skin and spinal cord, where 198 and 52 genes, respectively, displayed both altered methylation and expression profiles in MM lambs compared to unaffected control lambs (**Table 2, table S7**).

### Altered *ITGAD, PRSS53*, and *CALN1* expression reflects combined genomic and epigenomic control in MM

Our cross-tissue, multi-omics analysis suggests that altered expression of *ITGAD, PRSS53*, and *CALN1* may be jointly influenced by genomic variation (SNPs) and tissue-specific epigenomic regulation (DNA methylation), potentially contributing to MM pathogenesis. *ITGAD*, the top-associated gene within the Chr24 GWAS region, harbors hypermethylated CpGs and exhibits increased expression in spinal cord. *PRSS53* similarly displays consistent upregulation and hypermethylated CpGs in both muscle and skin. In contrast, *CALN1* exhibits hypomethylated CpG sites and reduced expression in the skin of MM animals.

### Functional annotation demonstrates key single- and multi-omics pathways underlying MM

Functional annotation of genes with GWAS signals highlighted several enriched biological processes, including cell-matrix and cell-cell adhesion, integrin-mediated signaling pathway connecting the extracellular matrix to the intracellular actin cytoskeleton, transcription, regulation of T cell activation, and amino acid biosynthesis.

In the muscle of MM lambs, DEGs were primarily enriched for pathways related to translation, immunoglobulin-mediated immune responses, MHC class II antigen assembly, type-I diabetes mellitus, cell adhesion molecules, and positive regulation of T-cell activation. Muscle DMGs showed enrichment for neural crest cell migration, focal adhesion, glutamatergic synapse, semaphorin–plexin signaling, neurogenesis, PI3K–Akt signaling, integrin-mediated signaling, and cell–matrix adhesion. In skin, DEGs were enriched for immune response, cell adhesion molecules, MHC class II antigen assembly, and positive regulation of T-cell activation. Skin DMGs were associated with glutamatergic synapse, long-term potentiation/depression, cytoskeletal organization in muscle cells, thiamine metabolism, and focal adhesion. In the spinal cord, DEGs were enriched for immune response, amino acid transport, NOD-like receptor signaling pathway, positive regulation of apoptotic process, inflammatory response and cytokine-cytokine receptor interaction. Spinal cord DMGs were enriched for phospholipid metabolism, regulation of small GTPase mediated signal transduction, cytoskeleton in muscle cells, long-term depression, mitosis, cilium biogenesis/degradation, neurogenesis, cell division and cell-cell adhesion (**table S8**).

Integration of multi-omics datasets further highlighted convergent biological processes. In muscle, overlaps among GWAS-DEG, GWAS-DMG, and DEG-DMG datasets were enriched for the AGE-RAGE signaling pathway in diabetic complications, positive regulation of keratinocyte migration, cell surface receptor signaling pathway, transcription initiation at RNA polymerase III promoter, and cell-cell adhesion. In skin, overlapping genes were enriched for transcription regulation and fibroblast proliferation. In the spinal cord, GWAS-DMG and DEG-DMG overlaps converged on genes involved in inflammatory responses, regulation of actin filament polymerization and protein stability, apoptotic regulation, immune activation, and modulation of cell communication. Additional enriched functions included regulation of transcription by RNA polymerase II, cell-matrix and cell-cell adhesion, and inner cell mass-associated proliferative programs, indicating that shared genetic, transcriptional, and epigenetic perturbations in MM lambs disproportionately affect pathways governing immune signaling, cytoskeletal dynamics, adhesion, and early developmental proliferation (**table S8**).

Overall, across tissues and omics layers, most observed biological processes converge on several major functional themes, including immune and inflammatory activation, adhesion and cytoskeletal remodeling, transcriptional regulation, neurodevelopmental and synaptic pathways, and metabolic and proliferative programs (**table S8**).

### Protein-Protein Interaction (PPI) networks highlight multi-omics regulatory hub genes of MM

To identify biologically central hub genes among GWAS hits, DEGs, and DMGs associated with MM, we constructed PPI networks and analyzed network connectedness and centrality using a unified gene set for each tissue. In muscle, 61 hub genes with a high degree of connectedness and centrality were prioritized among the genes previously associated with MM (**Fig. 5A**). Key examples include *ITGB6, DOCK4, COL1A1*, known for their role in cell adhesion, cytoskeleton regulation, migration and interaction with extracellular matrix (ECM) structural integrity. Additional hubs included *SHMT2* and *CBS*, which participate in folate/one-carbon metabolism, and *IGF2, PPARG, EP300*, which are widely implicated in early embryonic development and growth regulation. In skin, 60 highly connected hub genes were identified, including *ITGAL, ITGAV, ITGAM, ITGB8, and FBN,1* which play key roles in cell adhesion, cytoskeleton regulation, cell-cell and cell-ECM interactions and ECM structural integrity. Other prominent hubs included *GART* and *MAT1A*, associated with folate and one-carbon metabolism pathways, as well as developmental regulators such as *PPARG, EP300* and *ESR1* (**Fig. 5B**). The spinal cord network contained 57 hubs genes **(Fig. 5C)**, with enrichment for genes involved in adhesion and cytoskeletal organization, including *ITGAV, FBN1, DOCK4*. Additional hubs included *MTRR*, which functions in folate and one-carbon metabolism, and developmental regulators such as *CREBBP, PPARGC1A*, and *NOS1* (**Fig. 5C**).

**Figure 5.**
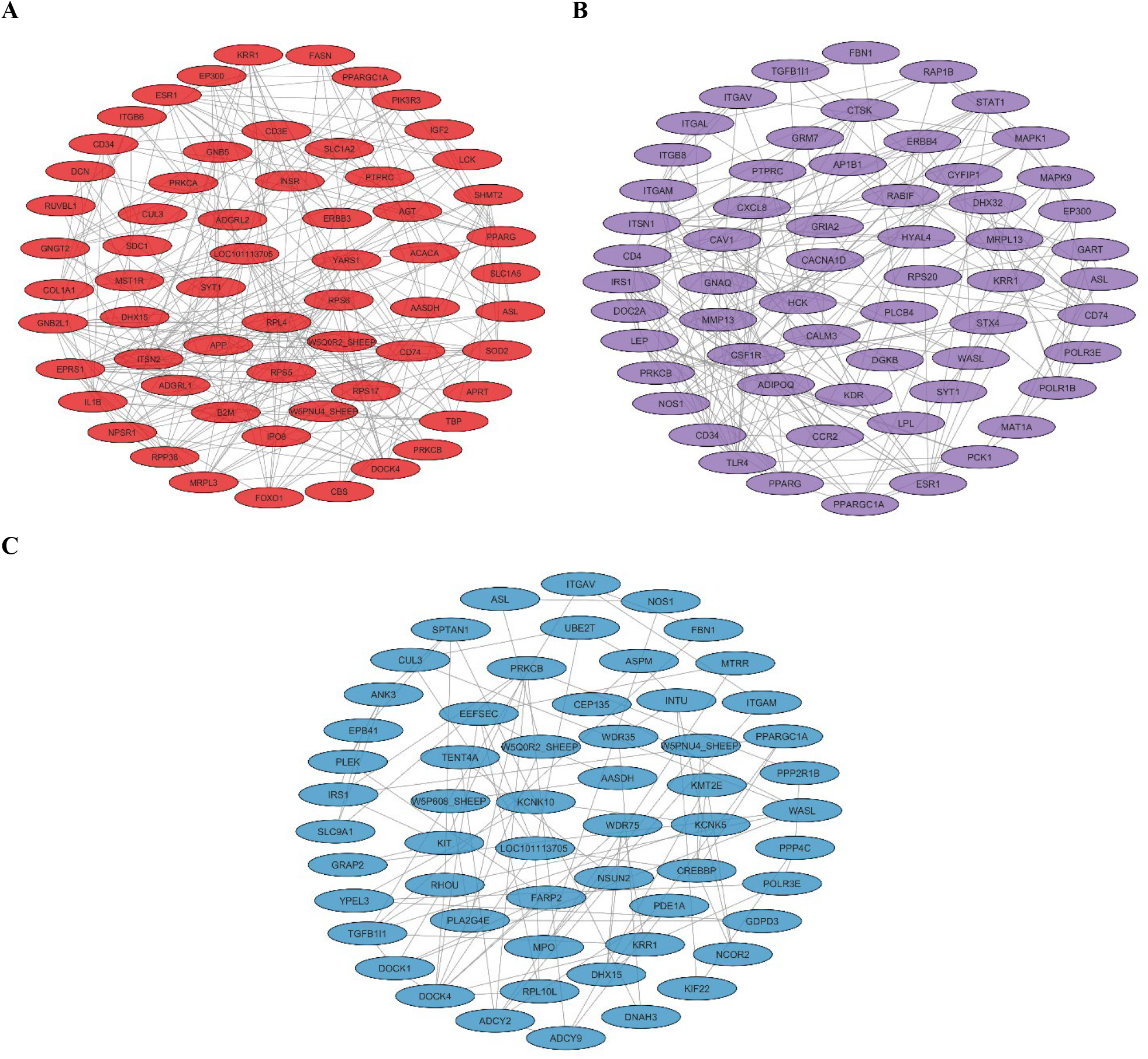
Hub genes associated with MM in muscle, skin, and spinal cord. **(A)** Muscle GWAS-DEG-DMG unified PPI network hub genes. **(B)** Skin GWAS-DEG-DMG unified PPI network hub genes. **(C)** Spinal cord GWAS-DEG-DMG unified PPI network hub genes.

Across tissues, several hub proteins were shared, indicating conserved regulatory roles. Eleven were common between muscle and skin (PPARGC1A, PPARG, EP300, SYT1, CD34, PTPRC, PRKCB, ESR1, CD74, KRR1, and ASL). Eleven proteins overlapped between muscle and spinal cord (PPARGC1A, PRKCB, W5PNU4, AASDH, CUL3, LOC101113705, DHX15, W5Q0R2, KRR1, and ASL). Twelve proteins were shared between skin and spinal cord (ITGAM, ITGAV, FBN1, TGFB1I1, PPARGC1A, POLR3E, PRKCB, NOS1, WASL, IRS1, KRR1, and ASL), highlighting recurrent network hubs that may represent core regulators of MM-related molecular pathways.

### Chr24 and Chr2 appear to be genomic hotspots for multi-omics signals underlying MM

The genomic distribution of GWAS signals, DEGs, and DMGs across muscle, skin, and spinal cord were visualized using karyotype plots to summarize multi-omics findings in MM lambs. The muscle karyotype plot (**Fig. 6**) highlights Chr24 as a major locus enriched for MM-associated genes overlapping across multiple omics layers. In addition, Chr2 and Chr3 showed notable enrichment in DMGs. A similar genomic pattern was observed in skin and spinal cord, where Chr24 again showed strong clustering of MM-associated genes **(Fig. S1, S2)**.

**Figure 6.**
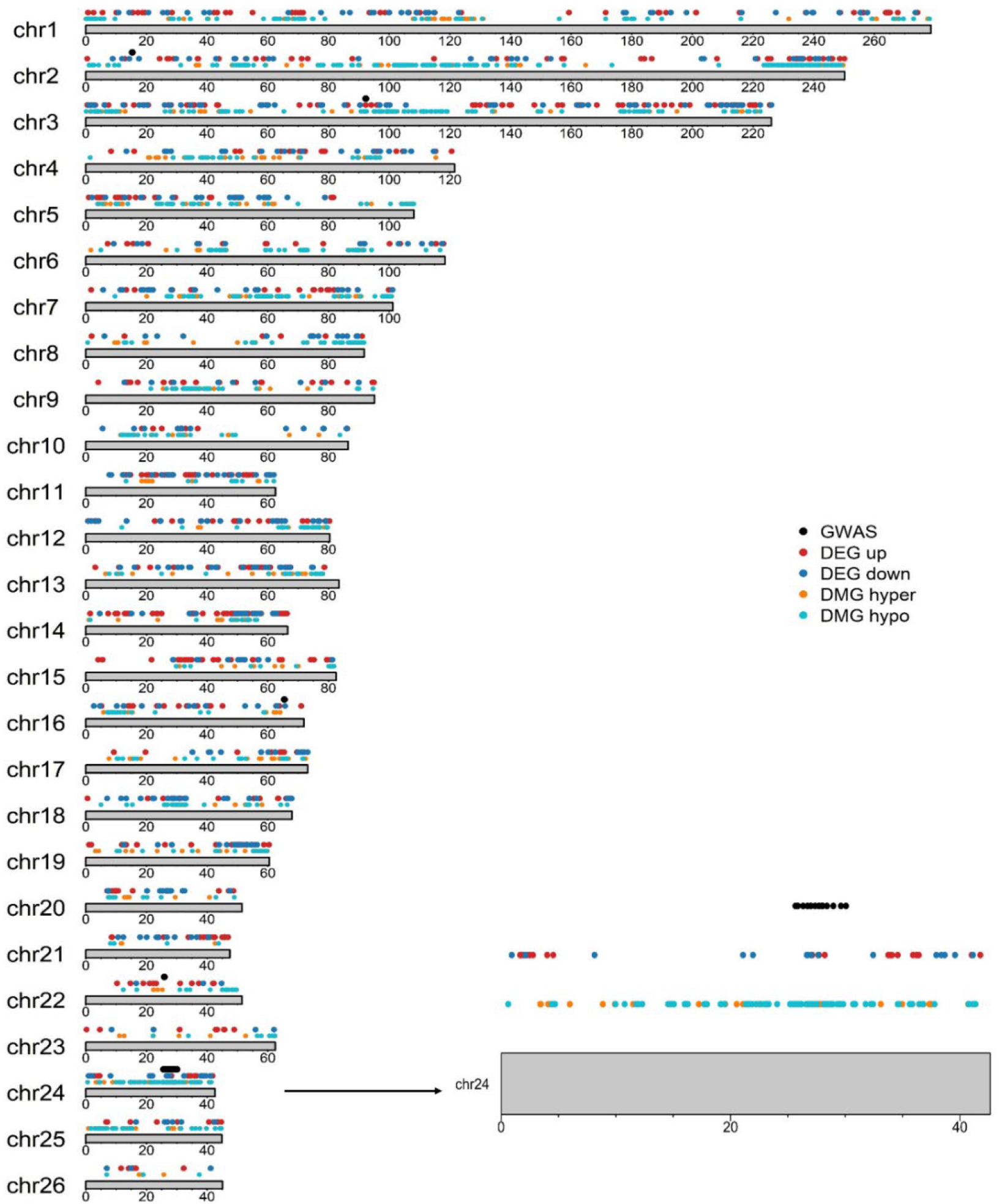
Genomic distribution to summarize GWAS, DEG, and DMG signals in the muscle tissue of MM lambs. The ARS-UI_Ramb_v3.0 reference genome assembly was used to determine marker locations. Chromosomes are organized by their actual length, which is indicated below each chromosome as checkpoints. Chr24 was further magnified (down-right) to show three-layer omics concordance on the 5 Mb significant interval identified by GWAS, DNA methylation, and gene expression analyses of MM.

## Discussion

In this study, using sheep as a large-animal NTD model, we investigated for the first time the heritability and multi-tissue molecular drivers of MM using integrated single- and multi-omics analyses in a mammalian system. These findings provide a rare window into the coordinated genetic, epigenetic, and transcriptional disruptions underlying MM naturally occurring in a sheep population. By integrating evidence across tissues central to neural tube closure and postnatal pathology, our results highlight both shared and tissue-specific pathways that converge on MM risk. Importantly, the identification of a heritable component alongside multi-omics signatures suggests that genetic predisposition interacts with downstream regulatory alterations to shape disease manifestation. Together, these insights establish a foundation for future functional studies and position this sheep population as a powerful comparative model for understanding MM biology and for developing translational intervention strategies.

We obtained moderate-to-high pedigree- and genome-based heritability estimates (0.68 and 0.42, respectively), indicating substantial genetic contribution to MM in this sheep population. Historically, only two small human twin studies have previously provided crude NTD heritability estimates (0.37–0.73) ^43,44^. Despite the limitations of twin-based designs, including the absence of dizygotic concordance in the California Twin Registry, our estimates closely parallel those in humans, supporting a similar gene-environment architecture for this NTD.

To investigate the genetic architecture of MM, we first conducted a GWAS that identified ∼5Mb genomic interval on Chr24 showing a strong association with MM frequency. The top associated gene, *ITGAD*, lies within a cluster of integrin genes including *ITGAM* and *ITGAX*. Other integrins (*ITGAL, ITGAV* and *ITGA4*) were also found to be associated with MM. Integrins encode the alpha subunit of heterodimeric cell surface glycoproteins that mediate cell-cell and cell-ECM adhesion, migration, and fusion ^45^. Many genes in this interval, including *ITGAM, ITGAX, GUSB, ASL, TYW1, CORO1A, CALN1, MTRR* and *TBX6*, have previously been linked to neurological traits, central nervous system and immune system disorders, as well as pairwise regulatory effects on each other in humans ^46–48^. Integrin signaling has been repeatedly implicated in neural tube closure through its role in cell adhesion and interactions with the WNT signaling pathway ^2,6^. Furthermore, *MTRR*, a gene involved in folate-dependent and one-carbon metabolism, is well-established as a contributor to spina bifida risk through its role in methionine metabolism and methyl group homeostasis ^49,50^.

To characterize epigenetic signatures associated with MM, we applied a multi-tissue DNA methylation analysis to identify lineage-specific and cross-lineage regulatory marks reflecting early developmental programming. Strikingly, Chr24 showed the highest proportion of DMCs in both muscle and spinal cord and remained among the most enriched chromosomes in skin (Fig. 2D, E, F). Cross-tissue comparisons revealed a subset of DMGs shared across multiple tissues. Because spinal cord, skin, and muscle arise from distinct embryonic lineages, the presence of shared, directionally consistent methylation changes suggests disruption of epigenetic programs established early in development. These cross-lineage signatures include genes such as *PAX3, NEK1, ITGA4, ITGAV, ITGAD, WDPCP, EPHA4, PPARGC1A, MTR, BMP4, AUTS2, FZD3, ASL, CALN1, KDM5B*, many of which are well-established for their roles in neural development, planar cell polarity, chromatic regulation, and morphogenetic processes of early embryonic development essential for neural tube formation ^47,51,52^. Notably, *PAX3, FZD3, BMP4*, and *MTR* have previously been associated with increased NTD risk, including MM ^6,26,53,54^.

RNA-seq profiling using the same multi-tissue design revealed widespread transcriptional dysregulation associated with MM. Neural tube closure in sheep occurs during approximately the third week of gestation, when germ-layer regulatory programs are still being established. Consequently, cross-tissue methylation and transcriptional alterations observed in MM lambs likely reflect early developmental perturbations rather than late, secondary responses. The intermediate PCA positioning of skin supports this interpretation by indicating partial retention of germ-layer lineage transcriptional signatures. Overall, these findings support a model in which epigenetic and transcriptional alterations established during germ-layer specification are propagated across descendant tissues and contribute to the molecular landscape of MM ^55,56^.

Consistent with this interpretation, we found 58 DEGs shared between muscle and skin, 23 DEGs between muscle and spinal cord, and 18 DEGs between skin and spinal cord, with 11 shared across all three tissues. These included several *cluster of differentiation* (CD) genes (*CD2, CD74, CD3D, CD3E, CD300E, CXCR6, CD27*, and *CD96*) known for their roles in neuroinflammation and cell adhesion. Additional genes such as *ESR1, FKBP5, TBX21*, and *IKZF3* play roles in transcriptional regulation, chromatin remodeling, and signaling pathways (*51*). Although CD molecules are widely studied in immune biology, many encode cell surface receptors and adhesion molecules, including integrins, that are crucial for cell-cell interaction, adhesion, cell migration, differentiation, and tissue morphogenesis during embryonic development ^57,58^. Previous studies have also demonstrated roles for CD proteins in germ-layer commitment in mesodermal and neural differentiation ^58,59^. Additionally, *ESR1* and several T-box transcription factors are known regulators of early embryonic cell fate decisions ^60,61^. Our findings, therefore, suggest that dysregulation of these regulatory genes across tissues may reflect coordinated developmental disruption in MM.

By integrating genomic, epigenomic, and transcriptomic layers, multi-omics and systems-biology frameworks reveal cross-layer regulatory cascades and causal variant-gene links that single-omics approaches miss, enabling more accurate and mechanistically grounded gene prioritization. Consistently, gene-level integration of GWAS signals with DMGs revealed several genes— including *ITGAD, ZG16, CRCP, GALNT17, KATNIP, AUTS2, GSG1L, CALN1, ASL, GUSB*, and *VKORC1L1*— that represent potential points of interaction between genetic variation and epigenetic regulation. Although none of these genes have previously been directly linked to MM, many have well-established roles in neural development, cell adhesion, and embryogenesis ^47,51,52^. For example, variants in *KATNIP (KIAA0556)* cause Joubert syndrome through disruption of ciliary scaffolding and microtubule dynamics ^62^. *AUTS2* influences cytoskeletal organization and WNT/β-catenin signaling during neuronal development ^63^, while *GALNT17* is associated with neurodevelopmental disorders, including Williams-Beuren Syndrome ^*64*^. *ASL* regulates arginine and nitric oxide metabolism important for germ-layer differentiation ^65–67^, and *GUSB* influences glycosaminoglycan metabolism involved in pluripotency, primitive ectoderm, and neuroectoderm commitment^*68*^. *CALN1* (*CABP8*) is a neural calcium-binding protein linked to schizophrenia ^69^, while the importance of Ca^+2^ gradient is well known for neural induction and neural/epidermal fate (*70*).

Integration of GWAS with transcriptional changes further highlighted genes such as *ITGAL, ITGAM, ITGAD, SPN, CALN1, CORO1A, LAT, DOC2A, PRRT2, HSD3B7, PSPH*, and *PRSS53*, many of which are involved in cell adhesion, immune signaling, neural function, and embryonic development (*47, 51, 52*). Among these, *CORO1A*, a cytoskeletal regulator, has previously been linked to neural signaling and NTD phenotypes in rat models ^70^. Other genes identified in this study provide additional mechanistic clues: *DCTPP1* regulates nucleotide metabolism, genomic stability, WNT signaling and DNA methylation ^71,72^, *PSPH* participates in serine-driven one-carbon metabolism ^73^, and *DOC2A* functions as a Ca^+2^ sensor implicated in neurodevelopmental disorders^74,75^. Integration between DMGs and DEGs hinted at transcriptional changes that may be dysregulated by altered DNA methylation. Genes such as *PAX3, NEK7, ITGAD, DOCK4, FBN2, KIF26A, WNT5B, EPHA2, NINJ2, NOS1, NXN, FKBP5, HOXD13*, and *HOXC13* are known for their cell adhesion, cytoskeleton regulation, immune response, neurological traits, and embryonic development associations ^9,47,51,52^. In particular, *PAX3, KIF26A*, and *WNT5B* have been previously implicated in the etiology of NTDs ^26,76,77^.

Pathway-level multi-omic analyses across tissues converged on biological processes including cell-cell and cell-matrix adhesion, cytoskeletal regulation, transcription, cell migration, immune activation, proliferation, apoptosis, and calcium mediated signaling pathways. These pathways overlap with canonical developmental pathways implicated in neural tube closure including PCP, WNT, Shh, folate-dependent one-carbon metabolism, contributing the biomechanics of neural fold elevation and fusion ^2,6,9–12,78–81^. Our results help bridge a key translational gap between mouse NTD genetics, in which more than 250 genes have been implicated, with the more limited human MM studies. The sheep model captures conserved developmental pathways while also revealing species-specific candidate genes likely shaped by differences in neurulation morphology and timing ^82–84^.

Network-based prioritization further identified central hub genes connecting multiple biological processes relevant to MM, including cell adhesion, motility, cytoskeletal regulation, embryonic development, and folate metabolism. These included integrin family members (*ITGAD, ITGAM, ITGB6, ITGAL, ITGAV*) and cytoskeletal regulators such as *DOCK4*, none of which have previously been strongly associated with MM risk. Additional hub genes also stand out as strong mechanistic MM candidates via folate dependent one-carbon metabolism (*SHMT2, CBS, GART, MAT1A*) ^85,86^, while *MTRR, IGF2*, and *EP300* are already linked to human MM risk ^27,53,87,88^. *MTRR*, is known to be part of folate dependent one-carbon metabolism, *IGF2* is an imprinted growth factor linked to placental and fetal growth, and *EP300* is a histone acetyltransferase required for early lineage specification and organogenesis. Another gene, *PPARγ*, is a key transcriptional regulator of trophoblast differentiation and early metabolic programming ^51^. Together, these hub genes represent biologically coherent candidates whose functions align with the developmental, metabolic, and structural processes implicated in MM. Shared hub genes across tissues, including *PPARγC1A, PRKCB, KRR1*, and *ASL* observed in all three, suggest a coordinated regulatory disruption reflected by both ectoderm- and mesoderm-derived tissues during early development.

Among individual genes supported by cross-tissue, multi-omics layers, *ITGAD, PRSS53*, and *CALN1* stand out as particularly strong candidates influenced by both genetic variation and DNA methylation changes that may jointly contribute to increased risk for MM. Muscle tissue exhibited the largest burden of DEGs and DMGs, followed by skin and spinal cord, suggesting a stronger contribution from mesoderm-derived tissues to MM pathology. Interestingly, karyotype visualizations consistently highlighted chromosome 24 as a dominant etiological hotspot for MM in this population. Furthermore, the comparative context further highlights a species-level resemblance: mouse models show a largely deterministic architecture, whereas human MM is far more heterogeneous and polygenic ^11,12^. Human MM risk reflects dispersed variation across folate/one-carbon metabolism, methylation, adhesion and cytoskeletal pathways, PCP and WNT/SHH signaling, and chromatin regulators, and is further amplified by strong gene-environment interactions, especially folate status, whereas comparable environmental effects in mice typically require genetically or epigenetically sensitized backgrounds. Our results reflect this complexity, revealing broad genetic and epigenetic heterogeneity across adhesion, cytoskeletal regulation, motility, folate/one-carbon metabolism, and chromatin-regulatory pathways. Collectively, these data support this sheep population as a highly representative large-animal NTD model, capturing the etiological architecture characteristic to human MM.

Together, these findings support an integrated mechanistic model for MM in which genetic, epigenetic, and transcriptional disruptions of early embryonic multi-layer signaling, differentiation, adhesion, and cytoskeletal regulators alter tissue stiffness and cell motility, destabilize focal adhesion-calcium feedback, and ultimately compromise neural fold formation and fusion at the affected site. Quantitatively, we prioritized genes supported by multi-omics layers across tissues derived from ectoderm and mesoderm that occupy critical biological functions. We present these as a candidate gene list for targeted human sequencing and functional follow-up. Importantly, we acknowledge that anatomical and gestational differences among sheep, mice, and humans may alter the relative contributions of specific genes involved in closure biomechanics. Accordingly, we frame our conclusions as mechanistically and biologically consistent and translationally prioritized, rather than as definitive causal claims. To accelerate validation, we recommend immediate testing of top candidates in human iPSC-derived neuroepithelial organoids and live-imaging assays to measure cell adhesion, protrusive behavior, and tissue force generation, complemented by targeted resequencing in well-phenotyped human MM cohorts to assess enrichment and gene-environment interactions. Such a coordinated pipeline would convert pathway-level sheep insights into human-actionable targets and clarify which species-specific signals reflect true human risk versus model biology. Overall, by integrating multi-tissue molecular layers, this study advances a testable, pathway-centric roadmap for MM genetics that both reconciles and extends mouse and limited human data.

This study was constrained by several factors. Lack of affected females and full-sib controls necessitated a male half-sib design to maintain statistical power and minimize confounding from sex imbalance and background genetic structure. Additionally, distinguishing primary etiologic signals from downstream responses remains challenging in any single-omics dataset. The multiomics, multi-tissue design therefore played a crucial role in prioritizing genes supported by multiple independent lines of evidence. Additional constraints included small sample size and high inbreeding inherent to the study population, which reduced GWAS power and mapping resolution. Expanded sampling, fine-mapping and functional validation studies are currently underway to address these limitations.

This study represents one of the most comprehensive multi-omics, multi-tissue systems-biology investigations aimed at identifying molecular drivers of a disease, and the first of its kind for NTDs. The prioritized genes identified here constitute candidates for future studies focused on validation in larger cohorts and functional testing through *in vitro* cell-culture and *in vivo* perturbation in mouse, sheep, and human cellular or organoid systems. These genes may also serve as potential biomarkers and targets for the development of diagnostic, prognostic, and therapeutic strategies. On the other hand, the epigenetic markers identified here represent the MM-associated signals across ectoderm- and mesoderm-derived tissues within a single generation. Although transgenerational epigenetic inheritance (TEI) has not yet been demonstrated for MM, the DMGs, including histone deacetylases, chromatin regulators, and folate/one-carbon metabolism genes, position TEI as a key direction for future lines of investigation in this population. Assessing heritable DNA-methylation and histone-acetylation signatures in the affected sheep lineage, together with embryo-based gene silencing assays (through genome-, epigenome- or RNA targeting), will be critical for establishing the functional relevance of these candidates. Prior work in the same species showed methionine-induced TEI persisting for up to five generations ^20^, suggesting that similar experimental designs could clarify the long-term incidence and recurrence of MM.

## Conclusions

By integrating quantitative genetics with multi-tissue genomic, epigenomic, and transcriptomic profiling, this study delivers the first systems-level dissection of MM and establishes sheep as a robust large-animal model for NTD research. We demonstrate that ovine MM is moderately to highly heritable and driven by coordinated dysregulation across developmental lineages, with convergent multi-omics signals, particularly at the chromosome 24 hotspot, highlighting biologically coherent pathways and a focused set of high-confidence candidate genes for functional validation through knockout and gene-silencing studies. These cross-layer signatures offer mechanistic insight into neural tube closure failure, embryonic loss, and compromised welfare in sheep, while nominating homologous human genes as compelling targets for sequencing and mechanistic studies in spina bifida. Together, these findings establish a foundational molecular atlas of MM and advance the path towards translational applications, including biomarker discovery, prevention strategies, and therapeutic intervention in spina bifida, leveraging a model system well suited for translation to human disease.

## Materials and Methods

### Ethics Approval & Study Reporting

All procedures involving animals were approved by the Institutional Animal Care and Use Committee of the University of Wisconsin-Madison (Animal protocols A006911-A04 and [A006893-A03](https://arrow.wisc.edu/arrow/sd/Rooms/DisplayPages/LayoutInitial?Container=com.webridge.entity.Entity%5bOID%5b40D06292C0BC11F0E092703DB0565000%5d%5d)). The authors followed ARRIVE guidelines throughout the study ^89^.

### Experimental population, tissue sampling and pedigree-based heritability estimation

Experimental study population was composed of a mixed-breed sheep flock maintained by the Bayliss Sheep Farm (Rushsylvania, OH, USA) between the years 2002-2024. Details of the population were previously described ^42^. Briefly, the population has been pedigreed since 2002 with lambing records including weight, birth date, sex, and litter size, and MM status (reformatted as 0-normal, 1-MM) since 2004. In 2025, a nucleus flock of 30 pregnant ewes and two rams was transferred to the Arlington Sheep Research Unit (University of Wisconsin-Madison, Madison, WI, USA) to maintain the population for further research.

Pedigree pruning, record standardizations and initial diagnosis were performed using various base R functions and ‘pedigreemm’ R package ^90,91^. For pedigree-based liability scale heritability (*h*^*2*^) estimates, we fitted a univariate threshold animal model for the binary trait using ‘MCMCglmm’ package in R ^92^. The model included an additive genetic random effect linked to the pedigree and an observation-level random effect to account for overdispersion. We analyzed MM with a Bayesian Monte Carlo Marcov Chain (MCMC) variance decomposition in which the observed phenotype arose from an underlying liability (*l*_i_). The model is outlined below:

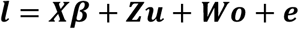

where, ***X*** and ***β*** are respectively the design matrix and vector of fixed effects including mean, ***Z*** and ***u*** are the design matrix and the vector of random additive genetic effects where 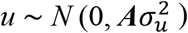,***w*** and ***o*** design matrix and vector observation-level random effect 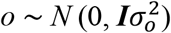,and 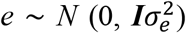, with 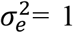 fixed under the probit link. ***A*** denoted here was a pedigree-based ‘expected’ numerator relationship matrix (NRM), while ***I*** was an identity matrix. Heritability on the liability scale was calculated as:

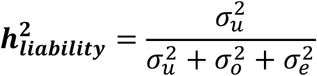

We used a moderately informative inverse-Wishart prior for both random effects (V = 0.5, v = 5) and ran 1,300,000 MCMC iterations, discarding the first 300,000 as burn-in and thinning every 1,000 iterations. Heritability was computed from each posterior sample, and the mean *h*^*2*^ was reported.

Among the population, 92 MM and 104 CT lambs (196 in total ) were sampled for semen, blood, or ear tissue between 2004 and 2025, randomly distributed across 21 generations. This sample set was used for genome-wide SNP genotyping to conduct GWAS, as described in the following section. Additionally, six male lambs born with MM in 2025, together with five of their half-sib CT pairs, were euthanized under the same conditions shortly after their birth. Muscle (mesoderm lineage), skin (mesoderm + non-neural ectoderm lineage) and spinal cord (neural ectoderm lineage) tissues from MM and CT lambs were collected for WGMS and RNA-Seq analyses described in the following sections. Tissues were sampled from the region close to the defect lesion in MM lambs, and from anatomically matching sites in CT lambs. Each sample was immediately divided into two portions: one was flash-frozen for WGMS, and the other was preserved in RNAlater for RNA-Seq. Both were stored in -80 ^°^C until further analysis.

### Genotyping, GWAS, and genomic heritability estimation

To investigate association between genomic variation and MM, we performed a GWAS using the 50K SNP genotypes of 196 animals (92 MM, 104 controls) and the binary response of MM. DNA extraction and genotyping with GGP Ovine 50K SNP array was conducted by a commercial provider following the manufacturer’s protocol (Neogen Corporation, MI, USA). Preceding further analysis, genotypes were imposed to quality control (QC) step to avoid possible genotyping errors and ensure higher imputation and GWAS accuracy. The ‘GenABEL’ R package was used to remove SNPs with low minor allele frequency <0.05, call rate <95%, and those located on sex chromosomes ^93^. Furthermore, samples with an individual call rate below 90%, identity by state (IBS) of >95%, and those with too high heterozygosity (False Discovery Rate < 1%) were removed from the data, which left 188 lambs (88 MM, 100 controls) and 41,389 SNPs for GWAS and genomic heritability estimation. Following QC, missing genotypes were imputed with expectation maximization (EM) algorithm ^94^.

For GWAS, a PCA-based approach (EIGENSTRAT) with a logistic regression model was used to effectively account for population structure and familial relatedness in ‘GenABEL’ R package ^93,95^. Briefly, a genomic relationship matrix (GRM) was calculated from SNP genotypes based on the model described below by ^96^:

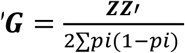

where, standardized by 2∑*pi*(1− *pi*) (i.e., allele frequencies), the **G** matrix resembles identity-by-state (IBD) based NRM from a pedigree but derived from identity-by-state (IBS) for ‘observed’ relationships. Subsequently, principal components (PCs) from the GRM were calculated, and the first three PCs were fitted into the GWAS model below, followed by an efficient score test for SNPs as described by ^95^. The model assumed a binomial response with a ‘logit’ link,

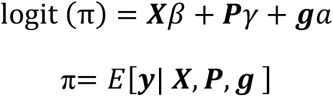

Where ***y*** denotes the 0/1 phenotype, ***X*** represents fixed covariates (e.g., intercept), **P** contains three principal components correcting for population structure, and **g** is the SNP genotypes coded for additive allele dosage effects (i.e., 0,1,2). A conservative Bonferroni-corrected genome-wide significance threshold was imposed with 0.05/41,389 (*p* < 1.20 x 10^-06^), followed by an average chromosome-wide representative threshold of (0.05/41,389) x 26 (*p* < 3.1 x 10^-05^) to account for test statistic inflation due to multiple testing. GWAS p-values are presented on Manhattan plots through -log10(p-value). Effect sizes were estimated as odds of allele dosage effects (i.e., odds of each additional copy of the effect allele).

For genomic heritability, the same threshold model applied for pedigree-based *h*^*2*^ above was used. Here, ***A*** was replaced by ‘observed’ relationships through ***G*** matrix and 88 MM cases vs. 100 controls were used with a moderately informative prior (V = 0.5, v = 3), 500,000 MCMC iterations of which 30,000 were discarded as burn-in and by thinning every 500 iterations.

Here, we estimated allelic dosage effects (i.e., additive effects) and reported them as absolute odds changes per single-effect allele. Therefore, any inferences for homozygous effect allele genotype should be made with twice the OR reported here, assuming additivity.

### Whole-genome methylation sequencing and high-resolution DMC analysis

To investigate the germ layer origins of MM and identify lineage-specific DNA methylation profiles underlying its pathogenesis, we performed WGMS for muscle, skin, and spinal cord tissues collected from regions adjacent to the affected area in both MM and CT lambs. Genomic DNA was extracted using the AllPrep DNA/RNA Mini Kit (QIAGEN, Hilden, Germany), following manufacturer’s protocol. WGMS was undertaken by the Genomics unit of the Roy J. Carver Biotechnology Center at the University of Illinois, Urbana-Champaign. Samples were sequenced with the Illumina NovaSeq X platform (Illumina, San Diego, CA, USA), generating on average 38x mean coverage of 148 bp length 1.18B paired-end reads per sample. ‘Fastq’ files were generated and demultiplexed using the ‘BCL Convert v4.3.16’ conversion software (Illumina).

An ASCII +33 quality score was used as Phred score for trimming adapters, and reads shorter than 40 bp were excluded from further analysis. Reads were then directionally aligned to the ‘ARS_UI_Ramb_3.0’ genome assembly through ‘DRAGEN v4.3.16’ (Illumina) software to obtain methylation calls. On average, 67.37 % of unique reads were mapped to the reference genome. One skin CT sample was eliminated due to failed QC. Methylation calls were extracted for CpG islands from ‘CX_reports’ through ‘awk’ software and transferred into R environment for differential methylation analysis ^91^.

To identify DMCs associated with MM, we compared CpG methylation profiles of muscle, skin, and spinal cord from MM vs. CT lambs at a single-base resolution. All DMC analyses were performed with ‘methylKit (v1.32.1)’ R package ^97^. Cytosines were filtered to exclude read counts ≤ 10, those on mtDNA, and methylation levels ≥ 99.9% from further analysis. This was followed by median normalization of the read coverages between samples. A beta-binomial hierarchical model with Bayesian dispersion shrinkage provided by ‘methylKit’ R package was used to detect CpG methylation count differences between MM and control groups. The dispersion corrected group means for each CpG were tested against the null hypothesis of ‘no difference’ by Wald’s F test, with a statistical model detailed by ^98^. The *p*-values were adjusted by a Benjamini–Hochberg false discovery rate (FDR) procedure to suppress the false positives due to multiple testing ^99^. Finally, two stringent significance thresholds were applied: only cytosines exhibiting a methylation difference of ≥ 20% between MM and CT groups and a q-value ≤ 0.05 were classified as DMCs. DMCs showing higher methylation in MM lambs were classified as hypermethylated, whereas those with lower methylation were classified as hypomethylated.

Following the detection of DMCs, the ‘ARS-UI_Ramb_v3.0’ was used with the R packages ‘genomation v1.38.0’ and ‘GenomicRanges v1.58.0’ to annotate DMCs with standard genic structures (i.e., promoter, intron, exon or intergenic) and the corresponding gene names, DMGs ^100,101^. DMGs were defined as genes with a DMC located within the promoter, gene bodies (exon, intron) or with a ± 20 kb distance to its transcription start site (TSS). Identified DMGs in each tissue were intersected to identify tissue-specific and coordinated changes across tissues from different embryonic layers linked to MM.

### RNA sequencing and differential gene expression analysis

To investigate lineage-specific and coordinated gene expression alterations associated with MM, we performed transcriptomic profiling by RNA sequencing using the same tissue samples used for WGMS. Total RNA was extracted using the AllPrep DNA/RNA Mini Kit (QIAGEN, Hilden, Germany), followed by DNase treatment. RNA sequencing was performed at the Genomics unit of the Roy J. Carver Biotechnology Center, University of Illinois at Urbana-Champaign. Accordingly, libraries were prepared using Watchmaker mRNA Library Prep Kit (Watchmaker Genomics, Boulder, CO, USA) with polyA selection. Subsequently, libraries were pooled, quantified by qPCR, and single-end sequenced on a 10B flow cell lane for 101 cycles using a NovaSeq X Plus with V1.0 sequencing kits to generate 100 bp reads. On average, 45.7M reads were generated per sample. ‘Fastq’ files were demultiplexed with the ‘BCL Convert v4.1.7’ software (Illumina). Following demultiplexing, an ASCII +33 quality score was used as Phred score for trimming and adapters were removed. High-quality reads were then aligned to the ‘ARS_UI_Ramb_3.0’ genome assembly through ‘STAR v2.7.5a’ aligner ^102^. Alignment of unique reads was, on average, 88.35% for samples. Read counts were obtained using the ‘Rsubread v2.20.0’ package on R environment ^103^.

To identify genes with altered expressions in muscle, skin, and spinal cord of MM lambs, we implemented differential expression analysis for each tissue using a logistic regression framework with a negative binomial distribution and Bayesian dispersion shrinkage implemented in ‘edgeR’ R package ^104^. Genes with ≥10 read counts in at least two CT and three MM lambs were kept for downstream analysis, while those not meeting this criterion were considered as having ‘no expression’ and discarded. Counts were normalized using Trimmed Mean of M-values (TMM) approach ^105^. Significance of the expression differences were tested using likelihood ratio test with a threshold of q <0.05 and FC ≥ 2, where p-values were corrected by FDR procedure described by ^99^. Additionally, PCA of the log transformed normalized read counts were also implemented to assess transcriptomic similarities and distances across tissues with alternate lineages. Overlaps between tissues were investigated to identify tissue-specific (i.e., lineage) and across tissue coordinated DEGs linked to MM.

### Functional annotation of the MM-linked genes

To identify pathway level associations with MM through single-omics and multi-omics overlaps, functional enrichment was performed independently for GWAS genes, DEGs, and DMGs in each tissue using DAVID bioinformatics software ^106^. For annotation, gene ontology (GO) and DAVID’s UP_KW biological processes and KEGG pathways were used and enrichment in terms of fold-change and adjusted Fisher’s exact test p-values were reported. For each tissue, pathways enriched in at least two omics layers were defined as convergent pathways, representing biological processes and KEGG pathways supported by multiple molecular signals.

### Multi-omics integration framework for gene and pathway prioritization

Complementary to the single-omics analyses, we followed a simple five-layer multi-omics integration framework for the molecular information layers obtained using the same population to further refine and prioritize genes and pathways linked to MM.

To capture tissue-specific multi-omics convergence at gene-level, we performed gene-level overlap analysis for each tissue. Intersections were computed between GWAS genes, DEGs, and DMGs, including all pairwise overlaps (GWAS-DMG, GWAS-DMG, DEG-DMG) and the three-way intersection (GWAS-DMG-DEG). Genes present in at least two omics layers were considered as multi-omics candidates, and those present in all three layers were designated high-confidence integrative hits.

To uncover pathway-level multi-omics convergence, for each tissue, pathways enriched in at least two omics layers were defined as convergent pathways, representing biological processes and KEGG pathways supported by multiple molecular signals associated with MM.

To highlight multi-omics regulatory hub genes and prioritize biologically central genes, we constructed tissue-specific Protein–Protein Interaction (PPI) networks using the union of GWAS, DEG, and DMG gene sets for each tissue. Networks were generated using STRING-DB *Ovis aries* (sheep) database through ‘Cytoscape v3.10.4’ ^107^. Genes with a high degree of connectedness (edges ≥ ∼10) and high centrality (∼ highest 10%) were prioritized for network analysis. These hub genes were further functionally annotated using DAVID software. Across-tissue overlaps were highlighted where observed.

Genome-wide distribution of GWAS hits, DEGs, and DMGs was visualized using tissue-specific karyotype plots. For each tissue, sheep chromosomal ideograms were annotated with actual physical length and separate tracks for GWAS-mapped genes, up- and down-regulated DEGs, and hyper- and hypomethylated DMGs. R packages ‘karyoploteR v1.32.0’ and ‘GenomicRanges v1.58.0’ were used to generate karyotype plots, annotate chromosomes, and set gene tracks. MM-linked genes from all three omics layers were plotted to highlight genomic regions on sheep chromosomes with convergent hotspots associated with MM.

After performing all analyses independently for muscle, skin, and spinal cord, we compared (i) multi-omics gene lists, (ii) convergent pathways, (iii) prioritized PPI network hubs, and (iv) genomic hotspots identified from karyotype plots. Genes and pathways shared among omics in each tissue and those shared among tissues were classified as core multi-omics signals, whereas those unique to a single tissue were considered tissue-specific candidates. Thus, gene and pathway sets resulting from these integrative processes were further designated as ‘candidate genes with strong multi-layer evidence pointing to the same biological mechanism.

## Acknowledgments

Authors thank Becky Johnson, Holly Hovanec, Kay Nelson, Michael Maroney, Jane Rieman, Jamie Reichert, and Todd Taylor for their assistance with the management of the sheep flock and sampling procedures. We also thank the UW-Madison Center for High Throughput Computing (CHTC) in the Department of Computer Sciences for providing computational resources to implement bioinformatics analysis.

## Author contributions

Conceptualization: MK, HK, BJI. Methodology: MK, HK, BJI. Data curation: MK, JNH. Project administration: MK, AAM, HK, BJI. Investigation: MK, AAM, JK, SPK, LAP, RSA. Formal Analysis: MK. Visualization: MK. Resources: MK, HK, BJI, TMG, JW, RHF. Supervision: HK, BJI. Funding Acquisition: BJI, RHF, JW, PASTURES Consortium.Writing—original draft: MK. Writing—review & editing: MK, AAM, JK, SPK, JNH, WF, LAP, RSA, JW, BC, RHF, HK, BJI.

## Data and materials availability

All data used in this study have been deposited in DRYAD and will be accessible through a link provided after the completion of the review process.

## Supplementary Materials

Supplementary materials of the study were provided in supplementary materials.pdf document and as Tables_S2-S9.xlsx.

