## Supplementary materials for "A Multi-Omics Atlas of Naturally Occurring Myelomeningocele in a Large-Animal Model Reveals Heritable Architecture"

Mehmet Kizilaslan *et al.*

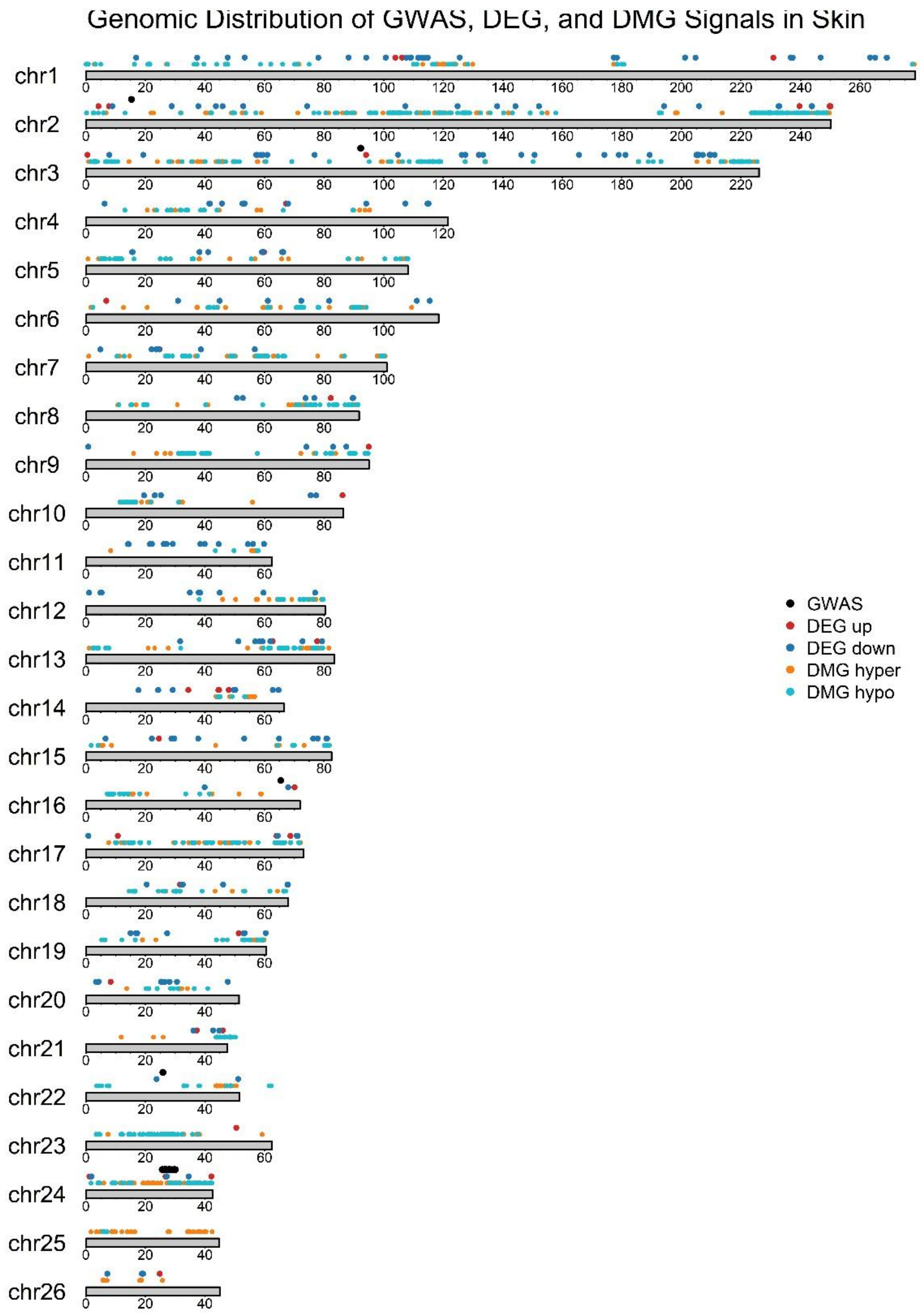


**Fig. S1.**

**Genomic distribution to summarize GWAS, DEG, and DMG signals in the skin tissue of MM lambs.** The ARS-UI_Ramb_v3.0 reference genome assembly was used to determine marker locations. Chromosomes are organized based on their actual length, which are designated below each chromosome as checkpoints.

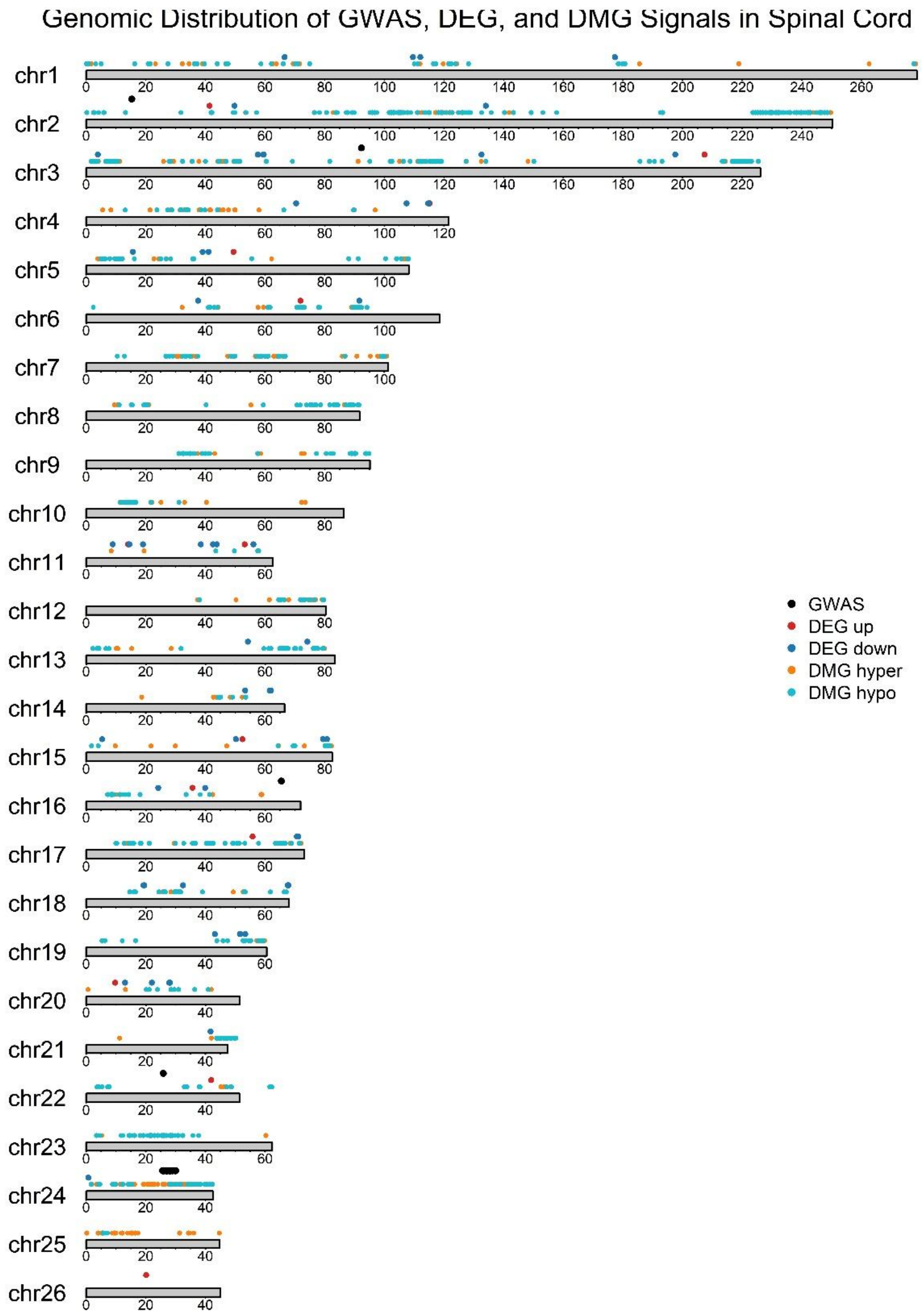


**Fig. S2.**

**Genomic distribution to summarize GWAS, DEG, and DMG signals in the spinal cord tissue of MM lambs.** The ARS-UI_Ramb_v3.0 reference genome assembly was used to determine marker locations. Chromosomes are organized based on their actual length, which are designated below each chromosome as checkpoints.

**Table S1.**

Descriptive information of genome-wide associated SNPs with MM in sheep.

| No | SNP | Chr | Position Oar_3.1 | Position ARS-UI_Ramb_v3.0 | OR (%) | MAF | Genes | Distance to gene (ARS-UI_Ramb_v3.0) | p-value |
|---|---|---|---|---|---|---|---|---|---|
| 1 | rs624251832 | 24 | 27,447,423 | 27,971,151 | 4.06 | 0.43 | *ITGAD*<br>*ITGAX*<br>*ITGAM* | Intron<br>~25 Kb<br>~50 Kb | $1.02 \times 10^{-10}$ |
| 2 | Chr24:27885694 | 24 | 27,885,694 | 28,408,832 | 4.57 | 0.40 | *GUSB*<br>*VCORC1L1*<br>*ASL*<br>*CRCP* | Intron<br>~10 Kb<br>~45 Kb<br>~60 Kb | $1.34 \times 10^{-10}$ |
| 3 | Chr24:25818842 | 24 | 25,818,842 | 26,328,350 | 3.70 | 0.35 | *SBK1*<br>*LAT*<br>*XPO6* | Intron<br>~ 45 Kb<br>~70 Kb | $5.51 \times 10^{-10}$ |
| 4 | Chr24:27505868 | 24 | 27,505,868 | 28,029,909 | 3.85 | 0.38 | *TGFB1I1*<br>*ITGAD* | Intron<br>~35 Kb | $1.07 \times 10^{-09}$ |
| 5 | rs624017013 | 24 | 27,179,284 | 27,702,624 | 3.57 | 0.37 | *ZNF646*<br>*ZNF668*<br>*PRSS5*<br>*STX4*<br>*VKORC1*<br>*PRSS8*<br>*PRSS36* | Intron<br>~5 Kb<br>~10 Kb<br>~15 Kb<br>~20 Kb<br>~40 Kb<br>~50 Kb | $1.49 \times 10^{-09}$ |
| 6 | oar3_OAR24_28402848 | 24 | 28,402,848 | 28,961,981 | 3.09 | 0.49 | *TYW1*<br>*CALN1* | Intron<br>~50 Kb | $9.30 \times 10^{-09}$ |
| 7 | Chr24:29062993 | 24 | 29,062,993 | 29,621,113 | 3.38 | 0.45 | *GALNT17*<br>*CALN1* | Exon<br>~50Kb | $1.54 \times 10^{-08}$ |
| 8 | oar3_OAR24_26864704 | 24 | 26,864,704 | 27,365,901 | 3.14 | 0.47 | *SRCAP*<br>*TMEM265*<br>*PHKG2*<br>*ZNF629* | Exon<br>~10 Kb<br>~20 Kb<br>~50 Kb | $1.82 \times 10^{-08}$ |
| 9 | rs635195147 | 24 | 29,509,020 | 30,065,757 | 3.15 | 0.47 | *GALNT17* | ~20 Kb | $2.28 \times 10^{-08}$ |
| 10 | rs632215928 | 24 | 25,348,193 | 25,867,032 | 3.09 | 0.45 | *GSG1L*<br>*KATNIP* | Intron<br>~20 Kb | $8.00 \times 10^{-08}$ |
| 11 | rs626715035 | 24 | 25,141,253 | 25,660,515 | 2.81 | 0.46 | *KATNIP* | Intron | $1.79 \times 10^{-07}$ |
| 12 | rs635327462 | 24 | 26,493,858 | 26,996,108 | 4.84 | 0.28 | *C24H16orf54*<br>*QPRT*<br>*SPN*<br>*ZG16*<br>*KIF22* | Exon<br>~10 Kb<br>~20 Kb<br>~20 Kb<br>~30 Kb | $3.48 \times 10^{-07}$ |
| 13 | Chr11:21716649 | 11 | 21,716,649 | NW_024599817.1: 25,843 | 2.77 | 0.30 | *TLCD3A*<br>*GEMIN4*<br>*LOC114116890* | Exon<br>~5 Kb<br>~20 Kb | $5.32 \times 10^{-07}$ |

| | | | | | | | | | |
|---|---|---|---|---|---|---|---|---|---|
| 14 | oar3_OAR24_26207296 | 24 | 26,207,296 | 26,712,782 | 2.87 | 0.44 | *TBX6*<br>*GDPD3*<br>*PPP4C*<br>*YPEL3*<br>*TLCD3B*<br>*CORO1A* | Intron<br>~5 Kb<br>~5Kb<br>~5Kb<br>~40 Kb<br>~60 Kb | 1.02 x $10^{-06}$ |
| 16 | rs614615566 | 16 | 65316981 | 65,509,391 | 2.41 | 0.47 | *ADCY2*<br>*CFAB90*<br>*FASTKD3*<br>*MTRR* | Intron<br>~30 Kb<br>~50 Kb<br>~60 Kb | 3.84E-05 |
| 17 | rs625161059 | 22 | 25290968 | 25,913,586 | 2.99 | 0.38 | *SORCS3* | ~300 Kb | 4.76E-05 |
| 18 | OAR2_15288514.1 | 2 | 15288514 | 15,351,940 | 8.73 | 0.09 | *LOC132659281* (lincRNA) | ~15 Kb | 9.91E-05 |
| 19 | oar3_OAR3_92258060 | 3 | 92258060 | 92,336,722 | 11.38 | 0.06 | *FAM136A*<br>*XDH* | ~5 Kb<br>~15 Kb | 1 x $10^{-04}$ |

**Notes:** SNPs are ordered based on significance. where the first SNP is the top associated based on p-value. Chr = chromosome; OR (%) = Odds ratio in percentage; MAF= Minor allele frequency. SNPs 16 and 17 were only identified in chromosome-wide significance level, while 18 and 19 were only suggestive.

**Table S2. (separate file)**
185 genes and predicted loci found in GWAS-linked 5Mb region associated to MM in sheep.

**Table S3. (separate file)**
Differentially methylated cytosines (DMCs) found in muscle, skin and spinal cord tissues linked to MM.

**Table S4. (separate file)**
Differentially methylated genes (DMGs) associated to DMCs found in muscle, skin and spinal cord tissues.

**Table S5. (separate file)**
Differentially expressed genes (DEGs) found in muscle, skin and spinal cord tissues linked to MM.

**Table S6. (separate file)**
GWAS-DMG overlapping genes in muscle, skin and spinal cord.

**Table S7. (separate file)**
DMG-DEG overlapping genes in muscle, skin and spinal cord.

**Table S8. (separate file)**
Functional annotation and enrichment analysis of single- and multi-omics associated genes in muscle, skin and spinal cord.

**Table S9. (separate file)**
PASTURES consortium members, affiliations and access information.
